# Assessment of dopaminergic neurotoxicity of CYP2E1-mediated trichloroethylene metabolites in *C. elegans*

**DOI:** 10.64898/2026.11.23.768671

**Authors:** Jocelyn Dicent, Meghan L. Bucher, Stephanie O. Nnaji, Carolina Duarte-Hospital, Jessica H. Hartman, Gary W. Miller

## Abstract

Trichloroethylene (TCE) is an industrial solvent associated with Parkinson’s Disease (PD). Studies have demonstrated that TCE undergoes extensive metabolism in humans, suggesting that TCE toxicity may be attributable to its metabolites. However, knowledge of how TCE metabolites contribute to neurotoxicity in PD remains limited. We exposed *Caenorhabditis elegans* (*C. elegans*) to the TCE metabolites trichloroacetic acid (TCA) and trichloroethanol (TCEOH) to determine if exposure disrupts dopamine-dependent behaviors, such as movement and egg retention, and impairs dopamine neuron viability. TCA and TCEOH were selected because we have demonstrated that they are either detected or likely present in mouse brain after a human-relevant inhalation exposure to TCE. Wild-type, GFP-expressing, and an α-synuclein-expressing strain were exposed to 5 mM TCEOH, 10 mM TCA, or control doses. Dopaminergic neuronal viability and movement was assessed following 5 days of exposure and revealed a lack of pronounced neurodegeneration or deficits in movement following either exposure. However, TCEOH exposure for 2 days resulted in changes in egg retention, suggesting that it may disrupt dopaminergic or serotonergic neurotransmission. Although *C. elegans*-based assays can identify changes in PD-relevant outcomes, this study suggests that TCA and TCEOH in this exposure paradigm display limited morphological or behavioral toxicity in *C. elegans*, warranting further investigation of these metabolites in additional *C. elegans* and mammalian studies.

## Introduction

Trichloroethylene, or TCE, is a lipophilic, volatile halocarbon used as an industrial degreaser and in dry-cleaning applications (Agency for Toxic Substances and Disease Registry [ATSDR] 2019). As a persistent environmental contaminant, TCE readily accumulates in groundwater and soil, resulting in exposure in the general population primarily through drinking water and vapor intrusion (Wu and Schaum 2000). TCE has been linked to a wide range of health outcomes, including increasing epidemiological and experimental evidence to suggest that exposure to TCE is associated with Parkinson’s disease (PD) (Guehl et al. 2003; Gash et al. 2008; Liu et al. 2010; Goldman et al. 2012; Keane et al. 2019; De Miranda et al. 2021; Ilieva et al. 2022; Goldman et al. 2023; Adamson et al. 2023). Although TCE exposure has been associated with PD, determining whether TCE metabolites contribute to neurotoxicity will improve our understanding of TCE, given that about 40–75% of TCE is metabolized upon exposure (Lash et al. 2000). Whether TCE metabolites act directly on dopaminergic neurons has received little to no investigation. For example, the primary metabolites of TCE from the CYP2E1 pathway of metabolism, trichloroacetic acid (TCA) and trichloroethanol (TCEOH), have been studied in the context of hepatotoxicity and renal toxicity, but are understudied in the context of neurodegeneration (Herren-Freund et al. 1987; Davis 1992; Poon et al. 2000; Mensing et al. 2002; Green et al. 2003; Fouad et al. 2013; Yaqoob et al. 2014; Ajeigbe and Oladokun 2024).

*Caenorhabditis elegans* (*C. elegans* or worms) are a non-parasitic, non-pathogenic, free-living soil nematode that has served as a model organism in numerous landmark discoveries in neuroscience and biology. *C. elegans* is a powerful model for studying gene-environment interactions, as it was the first complex organism to have its entire genome sequenced and shares substantial genetic similarities with humans (*C. elegans* Sequencing Consortium 1998). Its compact genome is amenable to transgenic manipulation to express human proteins, making it a relevant model for better understanding of PD-related neurodegeneration. The *C. elegans* model is well suited to studying dopaminergic neuron death, a hallmark of PD-related neurodegeneration, because *C. elegans* hermaphrodites have eight dopaminergic neurons that can be easily monitored for degeneration (Muralidhara and Hardege 2025). This study used two transgenic *C. elegans* strains. The first expresses green fluorescent protein (GFP) under the dopamine transporter promoter *dat-1p*, effectively labeling its dopaminergic neurons. The second expresses humanized α-synuclein and GFP under the *dat-1p* promoter, thereby labeling dopaminergic neurons and providing a Parkinson’s disease model (Cooper and Van Raamsdonk 2018). These strains are useful because they enable direct assessment of dopaminergic neuronal integrity following exposure to TCE metabolites.

Although TCE has numerous metabolites, we selected TCA and TCEOH as compounds of interest because they are primary TCE metabolites and there is limited information on their potential neurotoxicity. Additionally, we found that TCA can be detected and there is evidence supporting the presence of TCEOH in mouse brain after a human-relevant inhalation exposure (Dicent et al. 2026). Using *C. elegans* as a screening-amenable model, we explored the potential neurotoxic effects of these compounds by examining their effects on dopaminergic neuron integrity and behaviors influenced by dopaminergic signaling, such as movement and egg retention (Pribadi et al. 2023; Muralidhara and Hardege 2025). Our results inform future directions for studying the neurotoxicity of TCE metabolites and suggest that additional investigation of these compounds may be warranted.

## Materials and Methods

### Chemicals

TCA and TCEOH were purchased from Sigma-Aldrich (St. Louis, MO). NGM powder, pure agar powder, cholesterol, and the salts to make M9 buffer (monobasic potassium phosphate, dibasic sodium phosphate, sodium chloride, and magnesium sulfate), along with ampicillin and 5-Fluoro-2′-deoxyuridine (FUdR) for liquid culture, were purchased from Thermo Fisher (Waltham, MA). Additional chemicals used to make S-complete medium, including potassium citrate and trace metals mix powder, were purchased from Fisher Scientific and US Biological (Salem, MA), respectively. Potassium hydroxide, sodium hydroxide, 6 N hydrochloric acid solution, and Milli-Q water were purchased from Sigma-Aldrich. The hypochlorite solution used for synchronization was prepared using household bleach (Clorox, 8% sodium hypochlorite), Milli-Q water, and potassium hydroxide.

### C. elegans Husbandry

Standard culture methods, including the use of normal growth medium (NGM) plates or well plates for liquid culture, a culture temperature of 20 °C, and the OP50 strain of *E. coli* as a food source, were followed as described (Brenner, 1974). Use of different methods is mentioned in conjunction with the relevant assay. *C. elegans* strains used in this study include the wildtype N2 Bristol strain, BZ555 (*egIs1* [Pdat-1::GFP]), and UA44 (*baIn11* [Pdat-1::alpha-syn + Pdat-1::gfp]). The UA44 strain was provided as a gift from Dr. Catharine Rankin’s lab. All other strains were obtained from the Caenorhabditis Genetics Center, which is funded by the NIH Office of Research Infrastructure Programs (P40 OD010440).

All *C. elegans* (hereafter referred to as worms or *C. elegans*) that were used in these experiments were synchronized at the L1 stage of development, using a bleach/potassium hydroxide mixture. Worms cultured in liquid culture were maintained on an incubator shaker set to 120 rpm at 20 °C. Liquid cultures were maintained in S-complete media supplemented with ampicillin and 5-Fluoro-2′-deoxyuridine (FUdR) and fed using the UV-sensitive *E. coli* strain NEC937 following UV exposure, as previously described (Bradner et al. 2021).

### NGM-filled 6-well Plate Creation

NGM-filled 6-well plates were used for dosing for the egg retention assay and for video capture for the movement assays. Briefly, a 2.3% NGM solution was made using NGM powder (containing agar, sodium chloride, peptone, and cholesterol) and Milli-Q water. The NGM solution was autoclaved for 20 minutes at 121 °C (15 psi). After cooling to 60 °C, 1 M potassium diacidic phosphate solution (pH = 6), 1 M calcium chloride solution, and 1 M magnesium sulfate solution were added to the solution while heating and stirring, to a final concentration of 25 mM, 1 mM and 1 mM, respectively.

Low-evaporation 6-well cell culture plates were filled with 3 mL NGM per well in a laminar flow hood using serological pipettes. Once solidified, the plates were wrapped in parafilm and stored agar-side up at room temperature for 1 day to prevent condensation, then stored at 4 °C. Plates were warmed to room temperature before use.

### Synchronization

Gravid worms were collected from 6 cm NGM plates using Milli-Q water and transferred to 1.5-mL Eppendorf tubes. Tubes were centrifuged at 8,100 rcf for 1 minute, and the supernatant was discarded. 1 mL of bleach solution, prepared with 250 mM potassium hydroxide and 0.5% hypochlorite in Milli-Q water, was added to each tube. Tubes were placed on their sides and vortexed periodically until most of the worm bodies had dissolved, leaving the eggs. Tubes were then centrifuged again at 8,100 rcf for 1 minute, the supernatant was discarded, and the pellet was resuspended in Milli-Q water. This wash was repeated twice more before resuspending the eggs in 100 μL of Milli-Q water per tube. This was transferred to 1 mL of Milli-Q water (or more, depending on the egg concentration) in a 50-mL conical tube. Conical tubes were incubated in a shaking incubator at 120 rpm and 20 °C overnight to allow eggs to hatch and arrest at the L1 stage.

### pH Correction

pH is an important consideration in studies involving TCE metabolites, as several exhibit extreme pH values. *C. elegans* is resilient to a wide pH range, with one study demonstrating survival across pH 3.1 to 11.9 in the context of lethality (Khanna et al. 1997). Two other studies report transcriptional changes in C. elegans in response to alkaline or acidic environments (Hall et al. 2008; Cong et al. 2020). At baseline, TCA is highly acidic, with a pH of 1 at 81.7 g/L (500 mM) at 25 °C (Sigma-Aldrich). The literature suggests that pH correction of TCA with a sodium hydroxide solution is common in rodent models (Pereira 1996; DeAngelo et al. 1997; Plewa et al. 2002). To avoid confounding experimental results with potential pH effects, we similarly pH-corrected TCA doses before use.

10 M sodium hydroxide dissolved in M9 was used to pH-correct TCA doses. TCEOH, as an alcohol, did not require pH correction, but it did require DMSO for solubilization. All compounds were dissolved in M9 (or M9 with 0.3% DMSO for TCEOH), which has the greatest pH-buffering capacity of the solvents we tested; details are in the supplementary information (SI1). Based on additional pH testing to determine whether pH alone affected *C. elegans* experimental outcomes (SI1), TCA doses were pH-corrected to pH 7. pH strips and a pH meter were used to verify the pH of the doses before use.

### C. elegans Dosing

Synchronized populations were grown on 6-cm plates until the L4 stage of development, at which point worms were transferred to either NGM-filled 6-well plates or liquid culture to initiate dosing. Additional details about dosing medium can be found in the supplementary information (SI2). For NGM-filled 6-well plate dosing, treatment solution was pipetted directly onto the agar in small droplets and allowed to equilibrate for 30 minutes before adding worms. For liquid culture, the treatment was mixed with food, S-complete medium, and ampicillin, then distributed to the wells of a 48-well plate. Worms were dosed with either vehicle control (M9), solvent control (M9 and 0.3% DMSO), 10 mM TCA, or 5 mM TCEOH. These doses were selected after conducting a dose-response experiment assessing lethality, as described in the supplementary information (SI3). Doses were refreshed every 48 hours following the initial exposure. Due to the release of worms’ eggs in liquid culture and limited egg-laying period, dosing for the egg retention assay was carried out on NGM-filled 6-well plates, and worms were only dosed at a single time point at the L4 stage.

### C. elegans Egg Retention Assay

Egg retention assay was performed as previously described (Lee et al. 2024). Briefly, individual worms were placed in wells of a clear U-shaped 96-well plate, filled with 75 µL of a 20% bleach solution (in Milli-Q water) per well. The plate was observed in brightfield at the 4x and 10x objectives of an EVOS FL microscope. After the worm body was fully dissolved, the number of eggs was counted as a measure of eggs in utero. To avoid confounding the results due to the worms’ egg release in liquid culture, the egg retention assay dosing was carried out using 6-well plates. Synchronized populations were grown on 6 cm plates until the L4 stage of development, at which point worms were transferred to the 6-well plates to initiate dosing. This assay was selected because egg-laying / egg-retention is a dopamine-linked phenotype in *C. elegans* (Suo et al. 2002; McDonald et al. 2006; McMillen and Chew 2023; Pribadi et al. 2023)

### Fluorescent Microscopy Analysis

Worms were imaged on microscope slides with agar pads (2% pure agarose in Milli-Q water). Agar was pipetted onto a glass microscope slide, and a second slide was placed on top to flatten it into an even surface. Once solidified, the top slide was removed and a 30 μL drop of 5 mM levamisole was pipetted onto the center of the agar pad. Worms were placed in this drop before adding a coverslip to the slide. The slides were imaged using an APX100 microscope and GFP.

### Neuron size

Worms on microscope slides were imaged on an APX100 microscope at 20x magnification using GFP under consistent lamp-intensity settings. Z-stack images were acquired of the head region of each worm using a 1 μM step size and 27 steps. The optimal z-plane was determined by finding the plane where the most neurons in the head were in focus for imaging, and then imaging 13 steps above and below this plane. Exposure times were kept consistent across treatments within day and strain.

ImageJ and Fiji were used to create and apply a macro to consistently re-scale and pre-process images as necessary, such as generating a maximal intensity projection (MIP) image for all images and thresholding the images. CellProfiler was used to analyze neuron area size across different worm strains and treatments by identifying the neurons and recording the neuron area. To exclude any duplicate features during the analysis, the images were manually verified to confirm that the reported areas corresponded to the neurons in the image and that neurons from nearby worms were not counted. Details on the macros and the CellProfiler pipeline can be found in the supplementary information (SI4 and SI5).

### Dendrite Degeneration

Worms were imaged on microscope slides on an APX100 microscope at 60x magnification using GFP at consistent lamp-intensity settings. Z-stack images were acquired of the head region of each worm using a 1 μM step size and 27 steps. The optimal z-plane was determined by finding the plane where most of the dendrites in the head neurons were in focus for imaging, and then imaging 13 steps above and below this plane. Exposure times were kept consistent across treatments within day and strain.

ImageJ and Fiji were used to create and apply a macro to consistently re-scale and pre-process images as necessary, such as generating a maximal intensity projection (MIP) image for all images and adjusting the brightness of the photos.

Dendrite images were blindly scored with the open-access software Blinder, hosted by Solibyte Solutions (Durham, NC) (Cothren et al. 2018). The method was adapted from previous literature (Morton et al. 2023). Images were pre-processed using ImageJ and Fiji, using a macro that can be found in the supplementary information (SI4). Individual CEP neuron dendrites were scored in Blinder on a scale of 0 - 4, as follows:

- no visible damage or abnormalities
- blebs encompassing less than 50% of the dendrite
- blebs encompassing more than 50% of the dendrite
- breaks present with more than 50% of the dendrite remaining
- breaks present with less than 50% of the dendrite remaining

### Movement Assays

Worms were transferred from liquid culture to 1.5 mL Eppendorf tubes. Worms were washed with Milli-Q water to remove any bacteria on their bodies. Using NGM-filled 6-well plates, each set of worms was pipetted into a well. Once all liquid had completely evaporated or been absorbed, worms were observed after 5 minutes and again after 30 minutes. These time points were captured to emulate area-restricted searching (ARS) assays, in which worm behavior is typically assessed at two time points. Using an experimental design similar to an ARS assay lends itself to the expectation that worms will initially have more foraging behaviors when they are first placed on a plate, including increased speed, movement, and turns, that will later be reduced by 30 minutes once the worms have determined that there is no food on the plate. 30-second videos of the worms in 2 to 3 different areas of the well were captured using the FlyCap software and FLIR integrated imaging solutions camera. Videos were analyzed using the WormLab software.

### Statistical Analysis

Due to the multifactorial nature of many of the worm analyses, RStudio (Version 2025.09.2+418) was used to perform statistical testing. In general, worm analyses included an age factor (Day 1 vs. Day 2 or Day 1 vs. Day 3 vs. Day 5), a timing factor (5 minutes or 30 minutes), a strain factor (N2 vs. BZ555 vs. UA44), and a treatment factor (vehicle control vs. 10 mM TCA or solvent control vs. 5 mM TCEOH).

### Egg Retention Assay

Data were first normalized to the appropriate strain control (vehicle or solvent) as a percentage of control values before statistical testing. Statistical analysis was done using three-way ANOVA. Pairwise comparisons were conducted using estimated marginal means with Tukey’s correction. The analysis was conducted using the following R packages: tidyverse, janitor, readxl, multcomp, onewaytests, car, and emmeans (Wickham and RStudio 2023; Firke et al. 2024; Dag et al. 2025; Fox et al. 2026; Hothorn et al. 2026; Lenth et al. 2026; Wickham et al. 2026). Raw egg counts can be found in the supplementary information (SI6).

### Behavioral Assays

As movement had been assessed at multiple time points within a day, behavior was analyzed on day 5 of adulthood to simplify the analysis. Absolute peristaltic speed and wavelength were analyzed using the raw, untransformed data. Three-way ANOVAs with White’s heteroskedasticity correction were run to assess changes in these metrics across timing, strain, and treatment, followed by Sidak’s post-hoc test to adjust p-values for multiple comparisons.

Omega bending events were defined as periods in which the omega bend behavior occurred for at least 1 second (7 consecutive frames). Presence of omega bending behavior was analyzed using general logistic regression, followed by Sidak’s post-hoc test to adjust p-values for multiple comparisons.

High-angled bends were analyzed by first binning general bending behavior into low (< 60°), medium (60° - 90°), and high (≥ 90°) bending angles for simplicity. Three-way ANOVA with White’s heteroskedasticity correction was conducted in the high-bending angle bin to assess changes in posture across timing, strain, and treatment, followed by Sidak’s post hoc test to adjust p-values for multiple comparisons. Across all of the behavioral assays, a heteroscedasticity correction was applied to account for unequal variances.

All behavioral analyses were conducted using the following R packages: tidyverse, fs, beepr, afex, emmeans, multcomp, writexl, and broom (Wickham and RStudio 2023; Bååth and Dobbyn 2024; Ooms and McNamara 2025; Singmann et al. 2025; Hothorn et al. 2026; Lenth et al. 2026; Hester et al. 2026; Robinson et al. 2026).

### Neuron Area Assay

Data were first normalized to the appropriate strain control (vehicle or solvent control) as a percentage of control values before statistical testing. Statistical analysis was done initially using three-way ANOVAs, and then followed up by using two-way ANOVA within strains, with treatment and age as factors. A heteroscedasticity correction was applied to account for unequal variances. Analysis was conducted using the following R packages: tidyverse, janitor, car, afex, ggprism, and openxlsx (Wickham and RStudio 2023; Firke et al. 2024; Dawson 2025; Schauberger et al. 2025; Singmann et al. 2025; Fox et al. 2026)

### Dendrite Degeneration Assay

Statistical analysis was performed using pairwise chi-square tests to assess differences in the proportions of scores across day, treatment, and strain, while holding other factors constant. In an exploratory analysis, we used cumulative link mixed models (CLMMs) to account for repeated dendrite measurements within worms. This was followed up by pairwise comparisons using latent estimated marginal means with Tukey’s correction. The CLMM analysis can be found in the supplementary information (SI7).

## Results

To determine whether exposure to TCE metabolites results in dopaminergic neurotoxicity, worms were exposed to TCA and TCEOH, and egg retention, motor behavior, and dopaminergic neuronal morphology were evaluated (Figure 1). The N2 worm strain was included as a control strain for the BZ555 and UA44 worm strains, and the BZ555 strain was included as a control for the UA44 strain in the fluorescent reporter assays. The UA44 strain was selected for this study as an α-synuclein model of Parkinson’s disease in worms, given that it expresses human α-synuclein in the dopaminergic neurons, along with GFP. The UA44 strain has been shown to experience greater changes in neuronal health and movement as a result of treatment as compared to control strains (Ray et al. 2014; Gaeta et al. 2019; Starr et al. 2023; Promtang et al. 2024)

**Figure 1.**
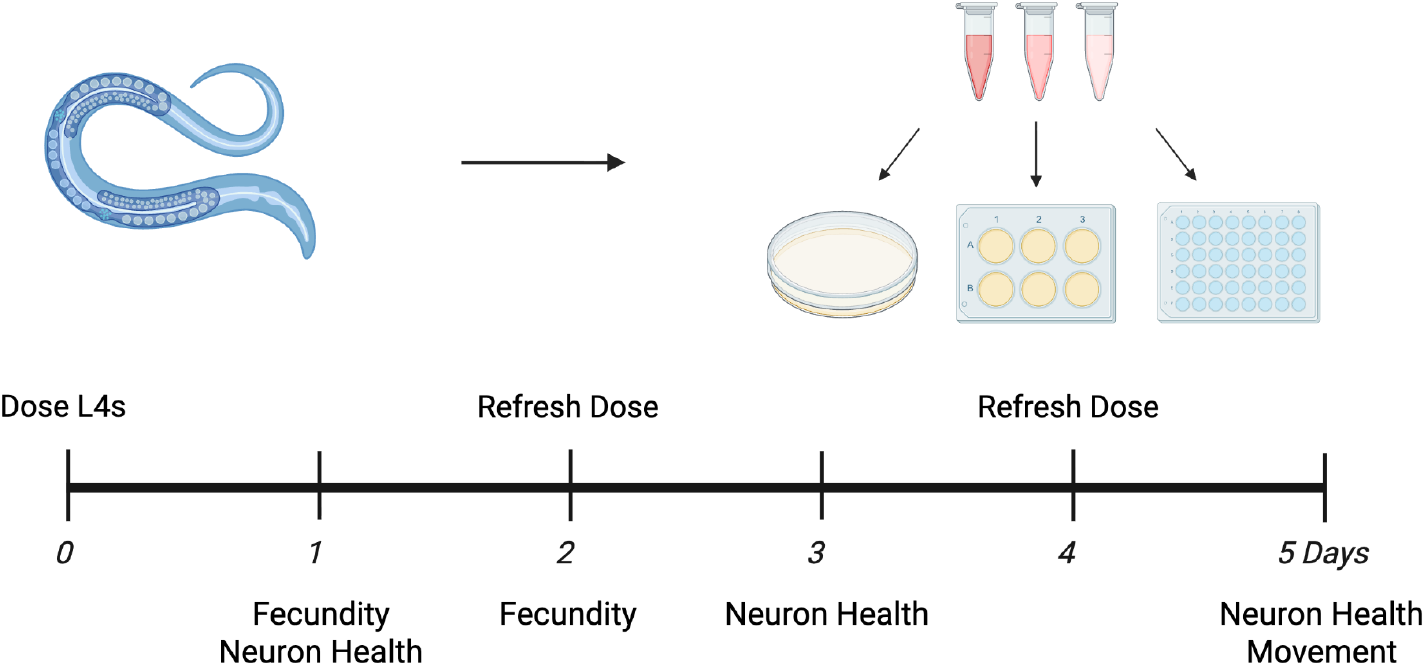
Dosing and experimental endpoints for *C. elegans*. Dosing was initiated using a synchronized population of L4s on either agar plates, agar-filled well plates, or 48-well plates for liquid culture. For experiments that went past Day 2 of adulthood, doses were refreshed every 48 hours. Depending on the experimental endpoints, worms were assessed at different developmental stages.

### Egg Retention

Egg retention for all strains was assessed at day 1 and day 2 of adulthood following exposure to TCA or TCEOH (Figure 2). Values were expressed as a percent of the control for that day and experiment, and then averaged across all experiments. Following TCA exposure, percent control values were largely consistent across strains, treatment, and days, with no statistically significant main effects or interactions (Figure 2A). In contrast, TCEOH exposure resulted in significant main effects of day, treatment, and strain, as well as significant day x treatment and day x strain interactions (Figure 2B). These interactions indicated that the treatment effect varied across developmental stages, precluding straightforward pairwise comparisons.

**Figure 2.**
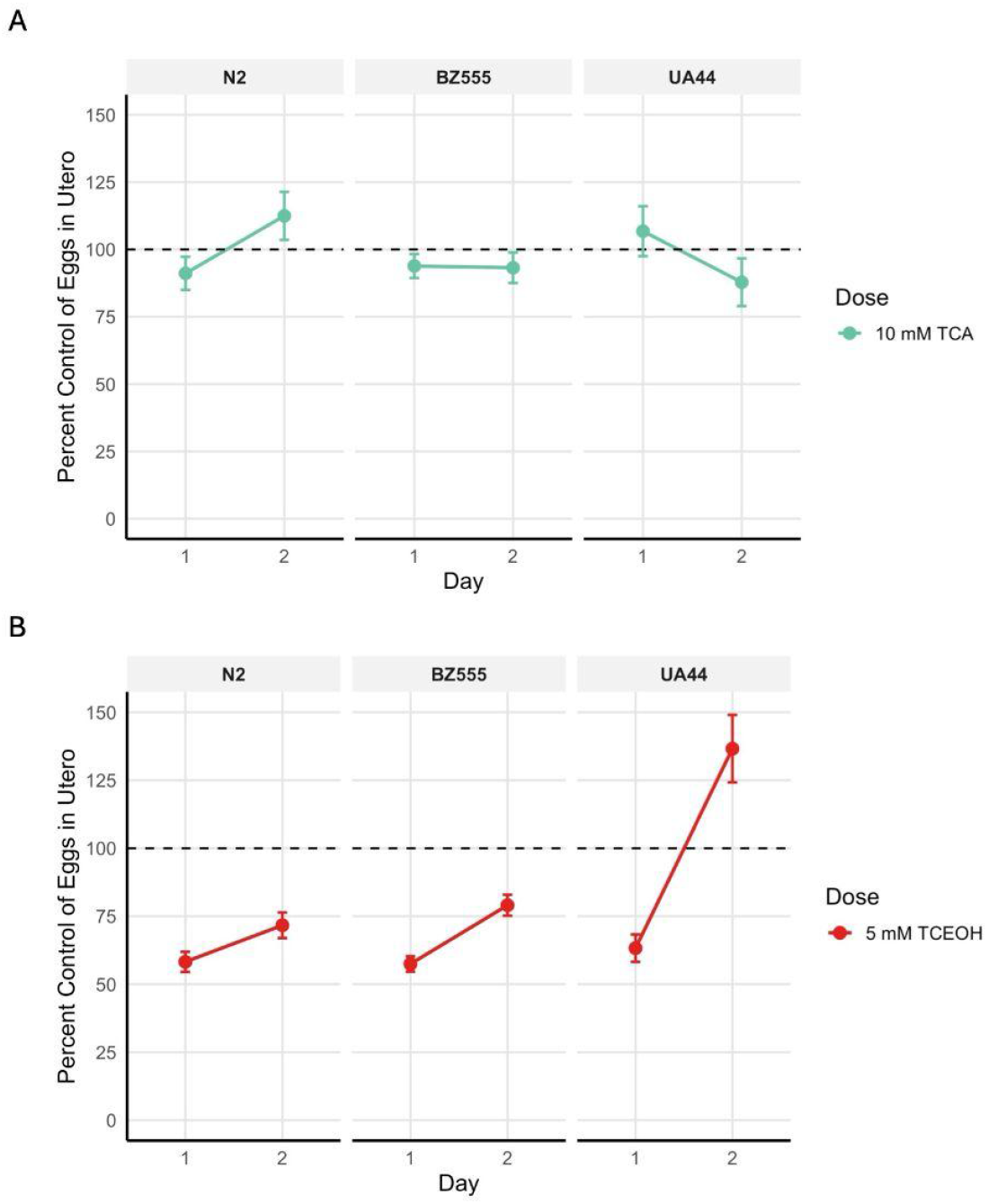
Percent control of eggs in utero across N2, BZ555, and UA44 strains across Day 1 and Day 2 of adulthood. **(A)** Trend graphs of the percent control of eggs in utero across strains and developmental stages after 10 mM TCA treatment. Three-way ANOVA indicated no significant differences among groups. **(B)** Trend graphs of the percent control of eggs in utero across strains and developmental stages after 5 mM TCEOH treatment. Three-way ANOVA indicated significant main effects of day, treatment, and strain, and significant day x treatment, day x strain, and day x dose x strain interactions (p < 0.05). The significant interactions prevented straightforward pairwise comparisons. TCA-treated groups are shown in green, and TCEOH-treated groups in red. The dashed line represents the respective control groups for each day and strain: vehicle control (M9) for the TCA group, and solvent control (DMSO in M9) for the TCEOH group. n = 35 - 40 worms per treatment, day, and strain across 4 experimental replicates.

Simple effects of TCEOH exposure were measured while holding other variables constant to account for the statistically significant interactions identified by the ANOVA (Tables 1, 2, and 3). First, we assessed the effect of day of adulthood while holding strain and dose constant (Table 1). Estimated marginal means showed that egg retention in TCEOH-exposed worms differed significantly between day 1 and day 2 of adulthood across all strains. Overall, all strains had decreased egg retention compared with the solvent control on day 1 of adulthood. The BZ555 and UA44 strains experienced a significant increase in egg retention by day 2 of adulthood, though the magnitude of this effect varied by strain. While the BZ555 strain showed a modest increase in egg retention that remained lower than the solvent control, the UA44 strain showed a severe increase, with egg retention higher than the solvent control.

**Table 1.** Effect of day within strain and dose on the TCEOH- and solvent-control-exposed groups in the egg retention assay. Given that for each strain and dose there was only one comparison (Day 1 vs. Day 2), p-values were not adjusted for multiple comparisons.

| **Strain** | **Dose** | **Day 1**<br>**EMM ± SE** | **Day 2**<br>**EMM ± SE** | **Difference**<br>**(Day 2 - Day 1)** | **t-ratio** | **p-value** |
|---|---|---|---|---|---|---|
| N2 | 5 mM<br>TCEOH | 58.20 ± 6.54 | 71.70 ± 6.54 | 13.50 | -1.45 | 0.15 |
| BZ555 | 5 mM<br>TCEOH | 57.40 ± 6.54 | 79.00 ± 6.54 | 21.6 | -2.35 | **0.02** |
| UA44 | 5 mM<br>TCEOH | 63.2 ± 6.54 | 136.6 ± 6.54 | 73.40 | -7.927 | **< 0.001** |

**Table 2.** Effect of strain within-day and dose on the TCEOH-exposed groups in the egg retention assay. Given that for each strain and dose there were three comparisons (N2 vs. BZ555, N2 vs. UA44, and BZ555 vs. UA44), p-values were adjusted for multiple comparisons using Tukey’s method.

| Comparison | Dose | Day | Estimate ± SE | t-ratio | Adjusted p-value |
|---|---|---|---|---|---|
| N2 - BZ555 | 5 mM TCEOH | 1 | 0.78 ± 9.25 | 0.08 | 1.00 |
|  |  | 2 | -7.38 ± 9.25 | -0.80 | 0.70 |
| N2 - UA44 | 5 mM TCEOH | 1 | -5.03 ± 9.25 | -0.54 | 0.85 |
|  |  | 2 | -64.94 ± 9.25 | -7.02 | **< 0.001** |
| BZ555 - UA44 | 5 mM TCEOH | 1 | -5.81 ± 9.25 | -0.63 | 0.80 |
|  |  | 2 | -57.56 ± 9.25 | -6.22 | **< 0.001** |

Next, we assessed the effect of strain while holding day and dose constant (Table 2). Overall, the strains responded similarly to the solvent control and TCEOH treatments. The exception is that for TCEOH-exposed worms on day 2 of adulthood, egg retention differed significantly between the UA44 strain and either the N2 or BZ555 strain. The response of the UA44 worms to TCEOH was greater than that of the N2 and BZ555 worms, which did not differ significantly from each other.

Finally, the effect of dose was assessed while holding strain and day constant (Table 3). Among day 1 adult worms, TCEOH exposure significantly decreased egg retention across all strains, with a slightly smaller effect in UA44 worms than in N2 and BZ555. Among day 2 adult worms, N2 and BZ555 worms exposed to TCEOH continued to show a significant decrease in egg retention compared with the solvent control, though egg retention increased compared with day 1 adults. In contrast, day 2 adult UA44 worms exposed to TCEOH showed a significant increase in egg retention compared with the solvent control, indicating a reversal of the effect in the TCEOH-exposed worms.

**Table 3.** Effect of dose within strain and day for the TCEOH- and solvent-control-exposed groups in the egg retention assay. Given that for each strain and dose there was only one comparison (Solvent Control vs. 5 mM TCEOH), p-values were not adjusted for multiple comparisons.

| Day | Strain | Solvent Control<br>EMM ± SE | 5 mM TCEOH<br>EMM ± SE | Difference | t-ratio | p-value |
|---|---|---|---|---|---|---|
| 1 | N2 | 100.0 ± 6.54 | 58.2 ± 6.54 | 41.8 | 4.52 | **< 0.001** |
|  | BZ555 | 100.0 ± 6.54 | 57.4 ± 6.54 | 42.6 | 4.60 | **< 0.001** |
|  | UA44 | 100.0 ± 6.54 | 63.2 ± 6.54 | 36.8 | 3.97 | **< 0.001** |
| 2 | N2 | 100.0 ± 6.54 | 71.7 ± 6.54 | 28.3 | 3.06 | **0.002** |
|  | BZ555 | 100.0 ± 6.54 | 79.0 ± 6.54 | 21.0 | 2.26 | **0.024** |
|  | UA44 | 100.0 ± 6.54 | 136.6 ± 6.54 | -36.6 | -3.96 | **< 0.001** |

### Movement

Behavioral outcomes of worm crawling behavior were assessed in the context of an area-restricted searching (ARS) assay. ARS is a commonly assayed behavior in *C. elegans*, as it indicates their foraging behavior (Hills et al. 2004; Muralidhara and Hardege 2025). Their foraging behavior is regulated by the dopaminergic and glutamatergic systems. In the presence of food, dopamine is released, promoting dwelling behavior in worms, whereas in its absence, the opposite occurs. *C. elegans* will turn more frequently and sharply as they begin foraging for food. As the time since their last encounter with food increases, they begin to turn less frequently and sharply.

Using WormLab, a number of movement metrics were recorded, of which we chose to focus on the following: peristaltic speed (um/s) [Absolute peristaltic track length/time], wavelength (um), proportion of worms exhibiting omega bend behavior, and changes in the proportion of worms with high-angled (> 90°) bends. We chose peristaltic speed to determine whether gross changes in worm movement would affect speed. Wavelength, omega bend behavior, and worms with high-angled bends were selected for their relevance to ARS assays, in that we would expect higher-angled bends, a greater number of turns, and a greater wavelength to occur as the worms begin to forage for food, and then decrease as time goes on after their last encounter with food.

Results of the three-way ANOVA on peristaltic speed indicated that the effect of TCA treatment on speed is not uniform, making it challenging to assign a single, consistent impact of the treatment across these conditions (Figure 3A). The only significant and biologically meaningful pairwise comparison was found in the N2 strain, where TCA initially induces greater speed in the worms, followed by a return to a speed similar to the 5-minute vehicle control in each strain. TCA-treated UA44 worms appear to follow a similar trend without meeting the significance threshold. Notably, most of the worms follow the expected trend of having faster-moving worms at the 5-minute time point, regardless of treatment, although the differences in speed are not statistically significant. The only exceptions to this are the control-treated N2 and UA44 worms, which exhibited the opposite trend. On average, the BZ555 and UA44 worms appear to be slower-moving than the N2 worms, with the UA44 worms having a significantly lower baseline average speed in both control and TCA-treated worms. Overall, the effect of TCA on speed was not consistent enough to indicate that there was an effect.

**Figure 3.**
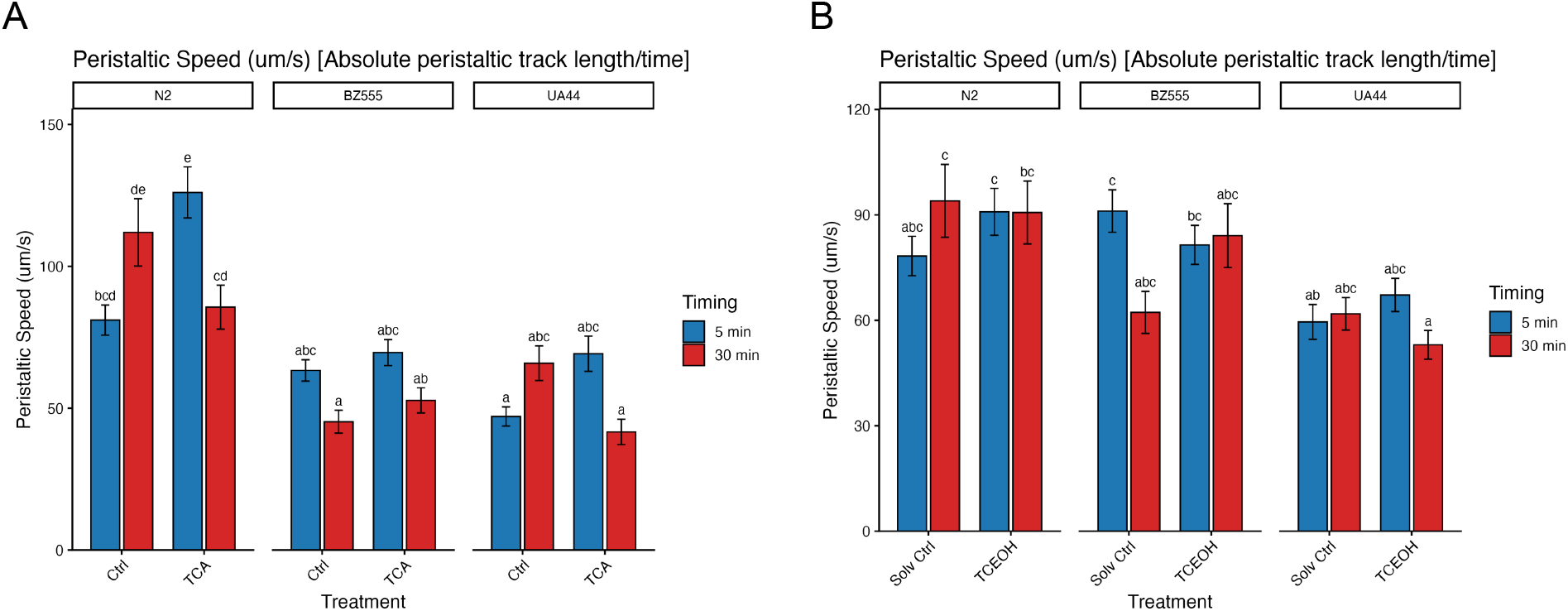
Peristaltic speed in N2, BZ555, and UA44 strains at day 5 of adulthood at 5 minutes and 30 minutes post-plating. Error bars represent ± SEM. Bars that share a letter are not significantly different from one another (Sidak, p < 0.05). **(A)** Peristaltic speed in vehicle control and TCA-exposed worms. Three-way ANOVA: timing (*F*(1, 935) = 6.45, *p* = 0.011), strain (*F*(2, 935) = 65.09, *p* < 0.001), timing × treatment (*F*(1, 935) = 30.86, *p* < 0.001), timing × treatment × strain (*F*(2, 935) = 9.15, *p* < 0.001). Non-significant effects omitted. **(B)** Peristaltic speed in solvent control and TCEOH-exposed worms. Three-way ANOVA: strain (*F*(2, 793) = 21.51, *p* = < 0.001), timing × treatment × strain (*F*(2, 793) = 4.56, *p* = 0.011) . Non-significant effects omitted. n = 42 - 114 worms per treatment, timing, and strain, across 3 experimental replicates.

Results of the three-way ANOVA on peristaltic speed indicated that there is no effect of TCEOH treatment on speed (Figure 3B). The differences noted in speed can be attributed to the significant main effect of strain or the interaction between strain, treatment, and timing. Similar to TCA, we note that UA44 worms appear to be slower-moving than N2 and BZ555 worms. Notably, most of the worms do not follow the expected trend of having faster-moving worms at the 5-minute time point, although the differences in speed are not statistically significant. While some pairwise comparisons were statistically significant, none were biologically meaningful. Overall, TCEOH does not appear to affect speed, though baseline strain differences in speed may indicate that the strains differ in overall locomotor performance.

Results of the three-way ANOVA on wavelength in worms treated with TCA and vehicle control indicated that the treatment effect depends on the strain, precluding straightforward conclusions about the treatment effect (Figure 4A). Unlike peristaltic speed, the worms appeared to have a more stable wavelength across strains, though subtle differences remained. In particular, the UA44 strain appeared to have a lower baseline wavelength compared to the other two strains. In addition, TCA treatment appears to increase wavelength at the 5-minute time point in the BZ555 strain, but not in the other strains. Notably, the trends in wavelength across time points are quite variable but are also not statistically significant. Overall, this suggests that any TCA treatment-related effects are subtle, which makes it difficult to define a single impact of TCA across these conditions.

**Figure 4.**
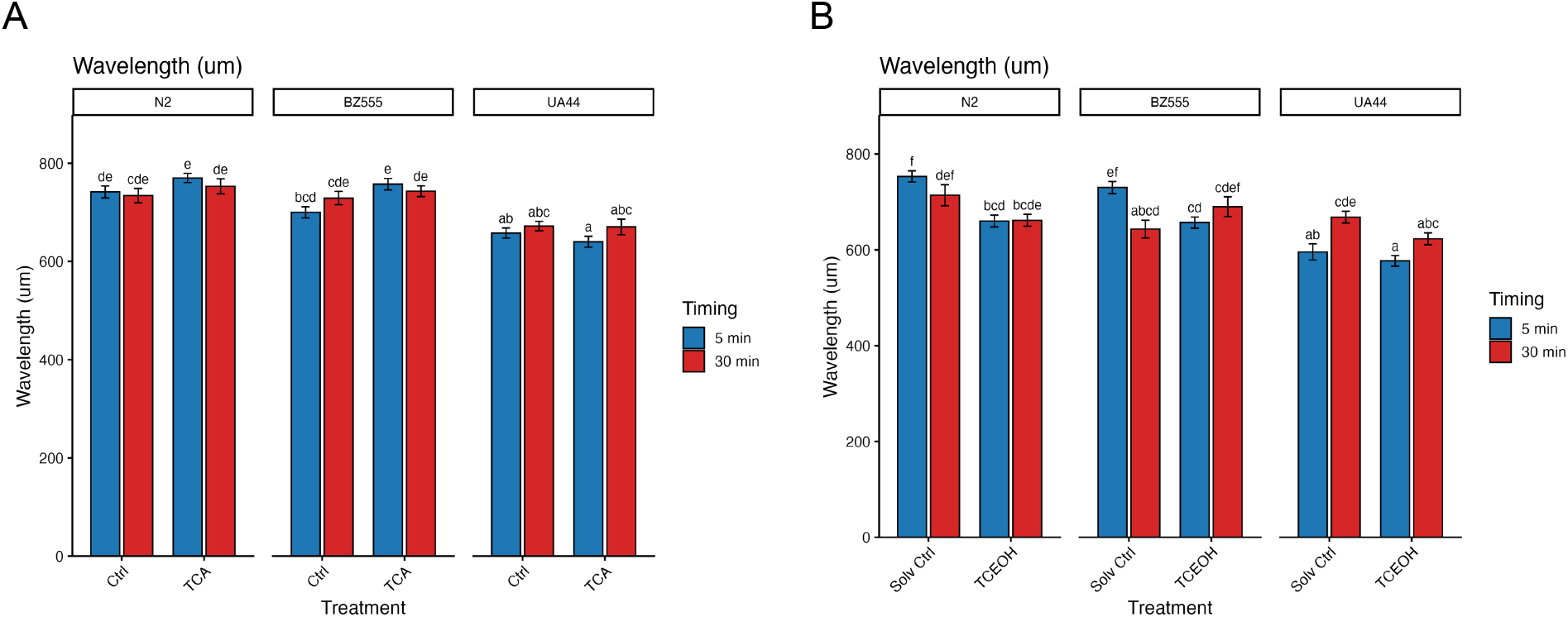
Wavelength in N2, BZ555, and UA44 strains at day 5 of adulthood at 5 minutes and 30 minutes post-plating. Error bars represent ± SEM. Bars that share a letter are not significantly different from one another (Sidak, p < 0.05). **(A)** Wavelength in vehicle control and TCA-exposed worms. Three-way ANOVA: treatment (*F*(1, 935) = 5.34, *p* = .021), strain (*F*(2, 935) = 60.83, *p* < 0.001), treatment × strain (*F*(2, 935) = 3.99, *p* = .019). Non-significant effects omitted. **(B)** Wavelength in solvent control and TCEOH-exposed worms. Three-way ANOVA: treatment (*F*(1, 793) = 19.91, *p* = < 0.001), strain (*F*(2, 793) = 34.38, *p* < 0.001), timing × treatment (*F*(1, 793) = 6.43, *p* = 0.011), timing × strain (*F*(2, 793) = 10.89, *p* < 0.001), treatment × strain (*F*(2, 793) = 3.60, *p* = 0.028), timing × treatment × strain (*F*(2, 793) = 6.28, *p* = 0.002). Non-significant effects omitted. n = 42 - 114 worms per treatment, timing, and strain, across 3 experimental replicates.

In comparison, the effect of TCEOH treatment on wavelength is dependent on both strain and timing (Figure 4B). Again, we note that the UA44 strain appeared to have a lower baseline wavelength compared to the other two strains. This difference appears to be driving the significant effects on treatment, as we observe that TCEOH-treated UA44 worms have a shorter wavelength at the 5-minute timepoint compared to other TCEOH-treated worms. Again, no clear pattern of effect emerges for TCEOH treatment, suggesting that its effect is likely subtle and largely controlled by strain and timing.

To assess the worm turning behavior, we calculated the number of worms that exhibited omega bending, defined as a worm conforming to an omega (Ω) shape (MBF Biosciences 2022). Omega bending can occur when a worm changes direction and is considered a high-angled turn, which is more common when worms exhibit foraging behavior. Since the data were binary, we assessed the probability that worms exhibited omega-bending behavior using logistic regression and plotted the proportion of worms exhibiting omega-bending in each group for clarity.

TCA-treated worms exhibited minimal differences in omega-bending behavior as compared to the controls, suggesting that TCA does not have an effect on bending behavior. The only significant finding of the logistic regression was that TCA-exposed N2 worms at the 30-minute time point had 80% lower odds of an omega bend compared to vehicle control N2 worms at the 5-minute time point, which is a finding that is not retained after pairwise comparisons (Figure 5A). Our analysis found no biologically meaningful pairwise differences, and we observed that the number of worms with omega bends was relatively similar across strains, time points, and treatments. The majority of the trends in omega bending were such that the 5-minute time point had a greater number of omega bends than the 30-minute time point within any treatment, which is consistent with what we would expect in worms that initially exhibit foraging behavior that is reduced over time. Overall, TCA treatment did not disrupt bending behavior.

**Figure 5.**
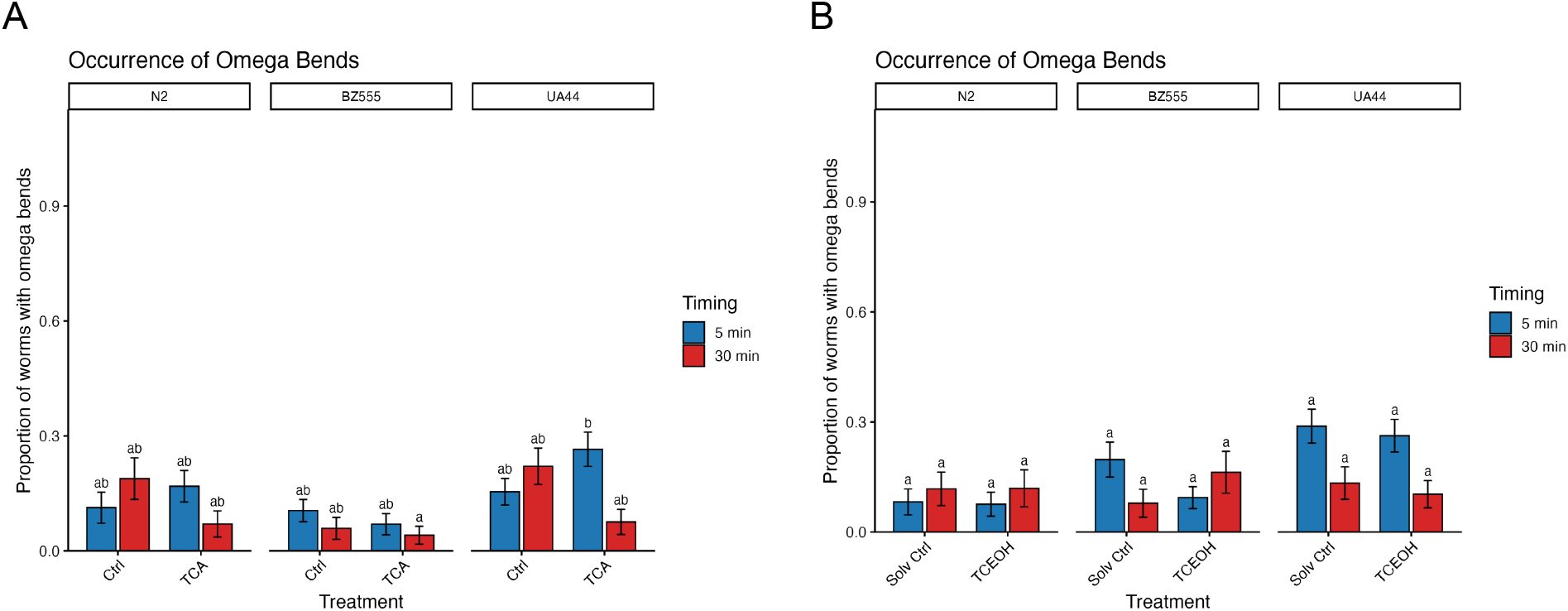
Proportion of worms with omega bends in N2, BZ555, and UA44 strains at Day 5 of adulthood at 5 minutes and 30 minutes post-plating. Error bars represent ± SEM. Bars that share a letter are not significantly different from one another (Sidak, p < 0.05). **(A)** Proportion of worms exhibiting omega bending behavior in vehicle control and TCA-exposed worms. Omega bending probability: logistic regression, TCA treatment × 30-minute timing (*z* = −1.99, *p* = .046, OR = 0.20). Non-significant effects omitted. (**B**) Proportion of worms exhibiting omega bending behavior in solvent control and TCEOH-exposed worms. Omega bending probability: logistic regression, UA44 strain (*z* = 2.92, *p* = .003, OR = 4.54). Non-significant effects omitted. n = 42 - 114 worms per treatment, timing, and strain, across 3 experimental replicates.

TCEOH-treated worms also exhibited minimal differences in omega-bending behavior as compared with the controls. The only significant logistic regression finding was that UA44 worms had 4.54 times higher odds of performing an omega bend than N2 worms (Figure 5B). This appears consistent with the plotted proportions, as UA44 omega-bend proportions are greater than N2 omega-bend proportions, though we observed no significant pairwise comparisons across any treatments, strains, or time points. Similarly, we find that the treatments vary greatly as to whether they have the expected behavior of initially exhibiting foraging behavior that declines over time. Overall, TCEOH appears to have no effect on the number of worms exhibiting omega-bend behavior.

We observed that treatment continued to have no effect when we considered any high-angled bending behavior in the worms, rather than only omega bending behavior (Figure 6). Although Figure 6 shows the distribution of high-angled bends across all frames of all worms per treatment, strain, and timing, we focused solely on high-angled bends (any bend > 90 °) for simplicity in the statistical analysis. Within this bending-angle category, we found no significant effect of TCA treatment on high-angle bends (Figure 6A). From the ANOVA, strain and timing had significant effects on high-angle bends; however, these effects are driven by comparisons between the TCA-treated N2 worms at the 30-minute time point and the vehicle-control-treated UA44 worms, and between the TCA-treated UA44 worms at the 5-minute time point and the vehicle-control-treated UA44 worms, which are not biologically meaningful comparisons. Overall, the relative amount of high-angled bending does not change much between time points, though there are fewer high-angled bends at the 30-minute time point than at the 5-minute time point.

**Figure 6.**
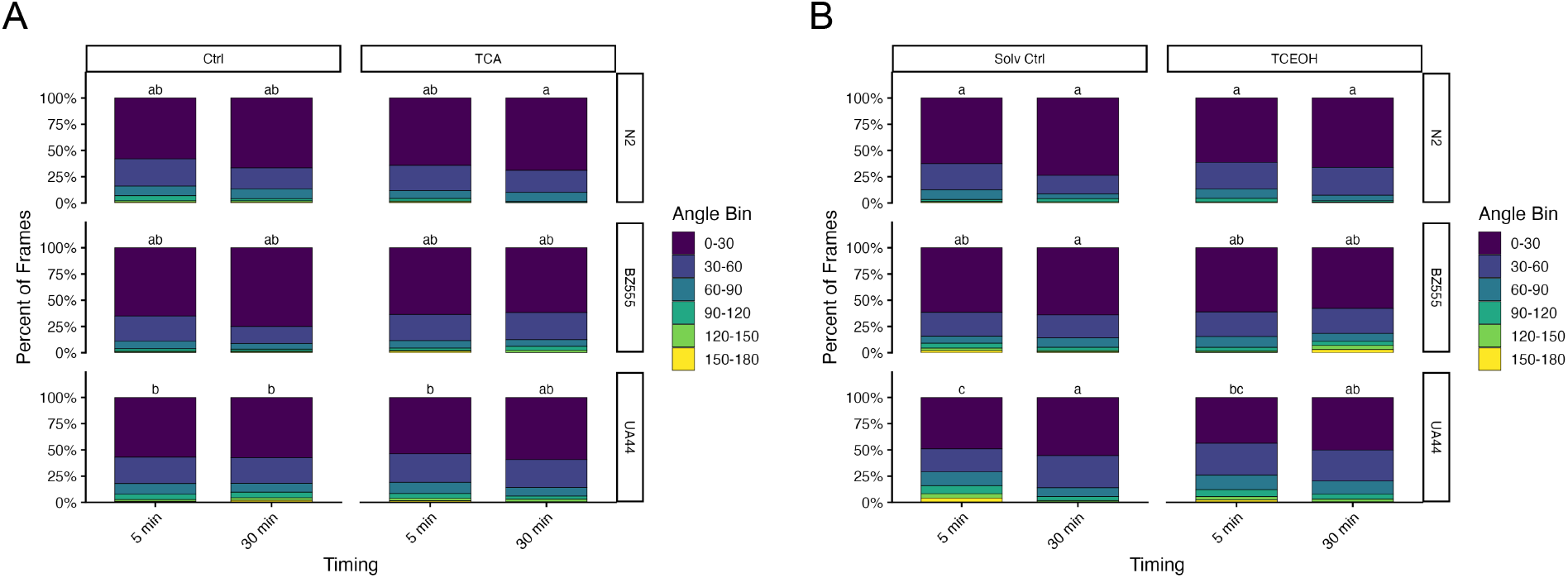
Distribution of bending angles across all worms and frames in N2, BZ555, and UA44 strains at Day 5 of adulthood at 5 minutes and 30 minutes post-plating. Bars that share a letter are not significantly different from one another for high-angled behavior (> 90°) (Sidak, p < 0.05). **(A)** Distribution of bending angles in vehicle control and TCA-exposed worms. Three-way ANOVA: strain (*F*(2, 12516) = 11.17, *p* < 0.001), timing (*F*(1, 12516) = 6.04, *p* = .014). Non-significant effects omitted. **(B)** Distribution of bending angles in solvent control and TCEOH-exposed worms. Three-way ANOVA: strain (*F*(2, 14676) = 21.66, *p* < 0.001), timing (*F*(1, 14676) = 21.26, *p* < 0.001), strain × timing (*F*(2, 14676) = 8.82, *p* < 0.001). Non-significant effects omitted. n = 42 - 114 worms per treatment, timing, and strain, across 3 experimental replicates.

For TCEOH-treated worms, the ANOVA results were the same as those of the TCA-treated worms (Figure 6B). Notably, the UA44 worms appear to have increased high-angled bending activity at baseline compared to the N2 and BZ555 worms. In addition, the TCEOH-treated UA44 worms have greater high-angled bending activity at the 5-minute time point as compared to the TCEOH-treated N2 worms at either time point. This suggests that UA44 may have increased movement relative to the N2 worms, attributable to strain differences rather than treatment. Although these comparisons are not significant, the general trend appears to be a decrease in high-angled bends at the 30-minute time point compared to the 5-minute time point. Overall, strain and timing, rather than treatment, influence high-angle bending behavior.

### Neuronal Morphology

Neuronal morphology was assessed at Day 1, Day 3, and Day 5 of adulthood to determine whether exposure to TCA or TCEOH impacts dopamine neuron health (Figure 7). Treatment with TCA or TCEOH did not significantly affect the cell body area as compared to the controls, indicating that there was no morphological effect on the dopamine cell bodies. Focusing solely on the neuronal cell bodies of TCA-treated worms, percent control values were largely consistent across strains, treatments, and ages, with no statistically significant main effects or interactions in the three-way ANOVA. In contrast, the three-way ANOVA for the TCEOH-treated group showed significant main effects of age, age:treatment, age:strain, and age:treatment:strain. Subsequent analysis by two-way ANOVAs (splitting the BZ555 and UA44 strains) resulted in a loss of the significant effects, suggesting that the significant differences identified by the three-way ANOVA were largely attributable to differences between strains rather than to treatment. TCEOH-treated worms also had percent control values that were largely consistent across strains, dose, and days. Thus, neither treatment significantly affected the cell body area and did not have a considerable impact on dopamine neuron health.

**Figure 7.**
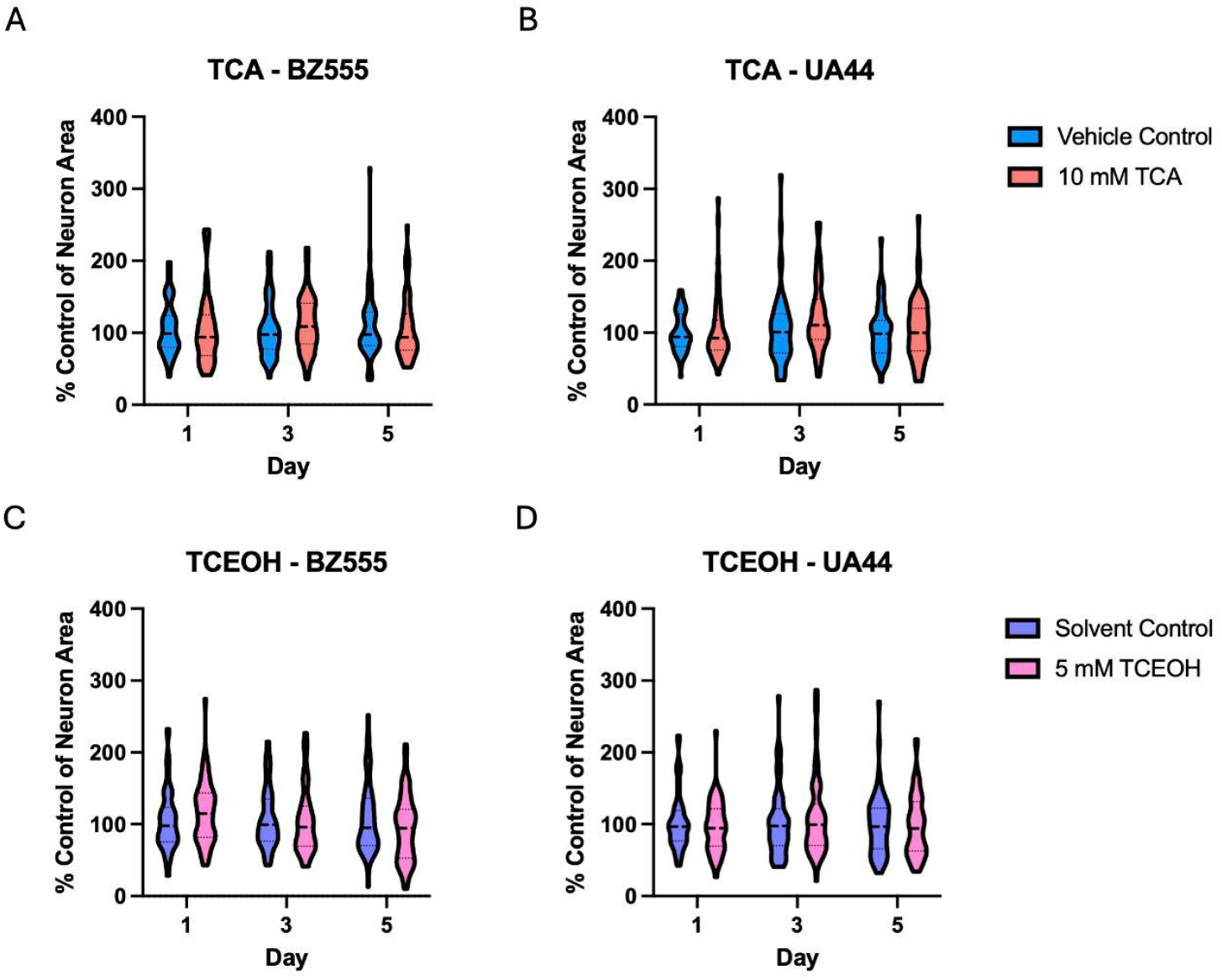
Percent control of neuron cell body areas in BZ555 and UA44 strains across Day 1, Day 3, and Day 5 of adulthood. Percent control of neuron cell body areas in the **(A)** BZ555 strain and **(B)** UA44 strain after exposure to 10 mM TCA. Three-way ANOVA revealed no significant effects (*p* > .14). Two-way ANOVA was used for each strain; no significant effects were detected in either strain (*p* > .06). Percent control of neuron cell body areas in the (**B**) BZ555 strain and **(D)** UA44 strain after exposure to 5 mM TCEOH. Three-way ANOVA: day (*F*(1, 2469) = 8.92, *p* = .003), day × dose (*F*(1, 2469) = 8.25, *p* = .004), day × strain (*F*(1, 2469) = 8.06, *p* = .005), day × dose × strain (*F*(1, 2469) = 7.50, *p* = .006). Non-significant effects omitted. Two-way ANOVA was used for each strain; no significant effects were detected in either strain (*p* > .09). n = 154–250 neuron cell bodies (from 58–62 worms) per treatment, age, and strain, across 3 experimental replicates.

Although there were no observed significant differences in cell body morphology, there were significant differences in the distribution of dendritic neurodegeneration scores (Figure 8). Nonetheless, these significant differences indicate that worm age and strain drive differences in dendritic neurodegeneration, and that treatment effect has a weak effect. Within the same day and treatment, the distribution of dendritic neurodegeneration scores differed significantly between BZ555 and UA44 worms that were treated with either TCA or vehicle control (p < 0.0001). Within the vehicle control-treated BZ555 worms, the distribution of dendritic neurodegeneration scores differed significantly between Day 1 and Day 5 adults (***χ***^2^ = 69.3966, p < 0.0001) and between Day 3 and Day 5 adults (***χ***^2^ = 55.5898, p < 0.0001). TCA-treated BZ555 worms differed significantly between Day 1 and Day 3 adults (***χ***^2^ = 24.3408, p < 0.0001) and between Day 1 and Day 5 adults (***χ***^2^ = 59.4249, p < 0.0001). Full results are in the supplementary information (SI6). Overall, this suggests that any effect of TCA treatment is weak and that worm age and strain drive differences in dendritic neurodegeneration.

**Figure 8.**
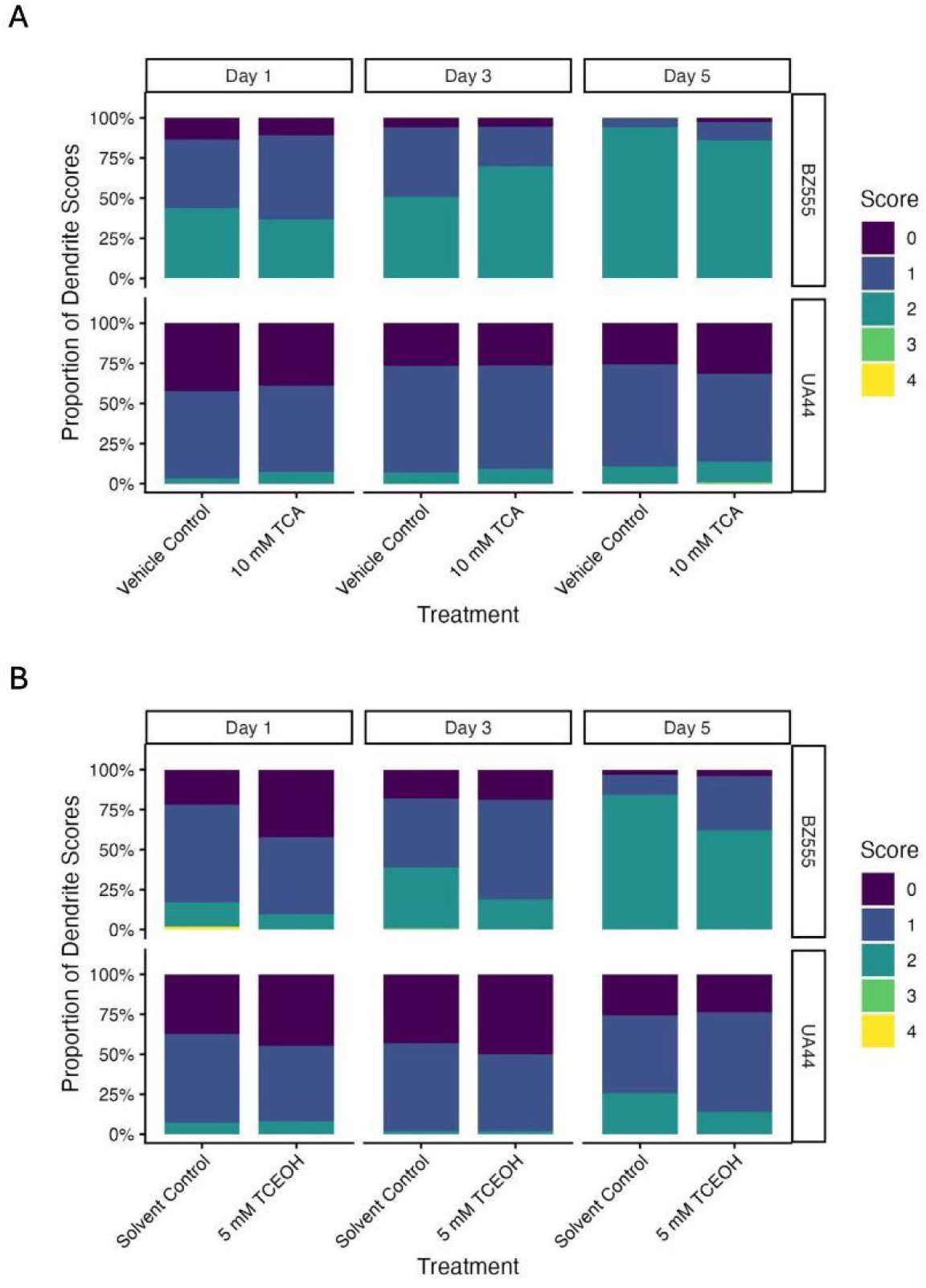
Distribution of dendrite degeneration scores (0 - 4) in BZ555 and UA44 strains across Day 1, Day 3, and Day 5 of adulthood.Proportion of dendrites assigned each degeneration score in BZ555 and UA44 strains after **(A)** exposure to vehicle control or 10 mM TCA or **(B)** exposure to solvent control or 5 mM TCEOH. Degeneration was scored on an ordinal scale of 0 - 4, with 0 indicating a healthy dendrite. Pairwise chi-squared analysis was run on the TCA and vehicle control-exposed worms and on the TCEOH- and solvent control-exposed worms separately using the Bonferroni correction to account for 24 pairwise comparisons. n = 108 - 126 dendrites (from 58 to 62 worms) per treatment, day, and strain, across 3 experimental replicates.

We observed similar results in TCEOH- and solvent control-treated worms. Within the same day and treatment, the distribution of dendritic neurodegeneration scores differed significantly between BZ555 and UA44 worms treated with either TCEOH or solvent control for Day 3 and Day 5 adults (p < 0.0001). Within the solvent control-treated BZ555 worms, the distribution of dendritic neurodegeneration scores differed significantly between Day 1 and Day 5 adults (***χ***^2^ = 111.6082, p < 0.0001) and between Day 3 and Day 5 adults (***χ***^2^ = 53.7607, p < 0.0001). TCEOH-treated BZ555 worms differed significantly between Day 1 and Day 5 adults (***χ***^2^ = 80.7378, p < 0.0001) and between Day 3 and Day 5 adults (***χ***^2^ = 46.1569, p < 0.0001). Solvent control-treated UA44 worms differed significantly between Day 3 and Day 5 adults (***χ***^2^ = 25.8177, p < 0.0001). Full results are in the supplementary information (SI6). Again, this suggests that any effect of TCEOH treatment is weak and that worm age and strain drive differences in dendrite neurodegeneration.

## Discussion

This study aimed to determine the effect of individual TCE metabolites on neurotoxicity-related outcomes in *C. elegans*. We chose the metabolites, TCA and TCEOH, because TCA was detected and TCEOH was likely present in the brains of mice exposed to TCE via a human-relevant inhalation dosing paradigm (Dicent et al. 2026). Furthermore, the volatility of TCE makes agar or liquid culture dosing unreliable. We focused on morphological characterization of dopaminergic neurons and dopamine-mediated functional outcomes in *C. elegans*.

All day 1 adult worms dosed with TCEOH had fewer eggs in utero, regardless of genotype. Meanwhile, the N2 and BZ555 strains continued to have fewer eggs in utero at day 2 of adulthood, though they appeared to recover partially compared to day 1 of adulthood. One possible explanation is that TCEOH may inhibit the C. elegans GρO ortholog GOA-1, mutations of which have been shown to cause hyperactive egg-laying (Bany et al. 2003). GOA-1 signaling inhibits presynaptic neurotransmitter release of serotonin and acetylcholine, so TCEOH may disrupt this inhibition, leading to extra neurotransmitter release and therefore additional egg laying.

Our laboratory has previously shown that worms with disrupted serotonergic signaling tend to hold their eggs in utero when treated with reserpine, a pharmacological inhibitor of catecholamine transporters (Bradner et al. 2021). In addition, worms with a mutation in the VMAT ortholog *cat-1*, which results in loss of the VMAT transporter, also hold their eggs in utero (Bradner et al. 2021). Notably, day 2 adult UA44 worms dosed with TCEOH appear to have the opposite effect compared with the other strains and days, retaining more eggs in a way that differs significantly from the control. Thus, TCEOH may impair neurotransmission or neurotransmission handling, resulting in the greater egg retention observed by Day 2 of adulthood in the UA44 strain. Overall, these findings suggest that TCEOH may affect dopaminergic or serotonergic neurotransmission, indicating that it may be a relevant metabolite to study in the context of monoamine neurotransmitter-related neurotoxicity. Additional tests may be necessary to determine exactly how TCEOH may impact neurotransmission or neurotransmission handling.

The movement analysis results indicate that behavioral changes were typically associated with either timing or strain. Effects of TCA or TCEOH treatment on behavior were mild, highly dependent on timing or strain, or too variable to detect. Regardless of treatment, baseline peristaltic speed and wavelength varied among worms, suggesting that locomotor function may differ by genotype. In particular, human α-synuclein appears to reduce speed and wavelength. Wavelength, as a measure of the sinusoidal movement of the worms, is expected to decrease with speed and suggests a change in the gait of the worms.

To further probe gait changes, we assessed the occurrence of omega-bend and high-angled-bend behaviors. These behaviors remained driven by strain or timing, with limited effect of TCA or TCEOH treatment. The UA44 worms had a higher probability of experiencing an omega bend than the N2 worms in the solvent control or TCEOH-treated worms. Although this trend was also observed in vehicle control or TCA-treated worms, it did not reach significance. A similar phenotype emerges when focusing on all types of high-angle bends; UA44 worms maintained high-angle conformations longer than N2 worms in the solvent control or TCEOH-treated worms. A similar trend occurs in vehicle-control and TCA-treated worms, though it does not reach statistical significance. The increased number of worms experiencing an omega bend or a high-angle conformation in the UA44 worms may be due to the presence of α-synuclein disrupting the dopaminergic system and leading to hyperactive turning, though additional analysis would be required to determine exactly which part of the dopaminergic system has been impacted to lead to this result (Muralidhara and Hardege 2025).

Our results indicate that changes in neuronal morphology depend strongly on worm strain and age, while the effect of TCA or TCEOH treatment on neuronal morphology is subtle. We found no significant differences in neuron cell body area across age, strain, or treatment in the TCA-treated group. Although the three-way ANOVA showed significant main effects and interactions in the TCEOH-treated group, these effects disappeared after conducting two-way ANOVAs within strains. Thus, differences in cell body area within the TCEOH-treated group were largely due to strain rather than treatment. Visually, UA44 worms appear to have slightly greater variability relative to the percent control of neuron cell body areas than BZ555 worms in TCEOH-treated worms.

Similarly, in the dendritic analysis, worm age and strain drove changes in neurodegeneration. Although there was a significant difference between the solvent control-treated and TCEOH-treated BZ555 worms across all ages, these significant results are lost to multiple-comparisons correction except for day 5 of adulthood. Notably, neither the UA44 nor BZ555 worms had a large proportion of severe neurodegeneration (scores of 3 or above). On average, BZ555 worms showed greater degeneration than UA44 worms, suggesting that α-synuclein may be neuroprotective against structural changes in neurons caused by both treatment-induced insults and age-related changes.

One limitation of this study is the lack of information on the internal dose of TCA or TCEOH, though we have evidence that both compounds are present in the worms’ systems (SI8). Without specific information about the internal dose, it is difficult to distinguish limited biological susceptibility from limited internal exposure. Low chemical uptake is a common limitation in *C. elegans* studies, and using higher concentrations to achieve higher uptake may be costly or limited by solubility (Hartman et al. 2021). In this study, we were limited by the solubility and the lethality of the compounds for our dosing. Additionally, mitochondrial biology may differ between *C. elegans* and mammals, suggesting that worms may generally be more resilient to complex I inhibitors. There is evidence that TCEOH is a mitochondrial toxicant, and work from others on classical PD-associated mitochondrial toxicants, like rotenone, have shown limited toxicity in *C. elegans* (Gonzalez-Hunt et al. 2021). While there is no evidence that TCA is a mitochondrial toxicant, future studies could test this, since its structural similarity to TCEOH and TCE makes this a possibility.

Our findings support a model in which TCE metabolites may affect the nervous system. Understanding the toxicity of TCE metabolites may help us better understand potential mechanisms of TCE neurotoxicity, though additional studies are needed to establish whether metabolites can mediate the neurotoxic effects of TCE itself. TCEOH, in particular, may disrupt dopaminergic neurotransmitter signaling without causing structural changes in dopaminergic neurons at a sublethal dose. These results are specific to the concentrations, exposure durations, and endpoints examined and suggest early, nuanced neurodegenerative effects from exposure to TCEOH. Future studies, either in *C. elegans* or in a mammalian model, assessing chronic exposure and targeted mechanisms will help to determine the long-term consequences of TCE metabolite exposure and their relevance to human health.

## Funding

The author(s) declare that financial support was received for the research, authorship, and/or publication of this article. JD received support from the National Institute of Environmental Health Sciences T32 ES007322. SN received support from the National Institute of Health grant R01ES023839. MB received support from the National Institute of Environmental Health Sciences T32 ES007322 and Parkinson’s Foundation Postdoctoral Fellowship PF-PRF-933478. GM was supported by the National Institute of Health grants RF1AG066107, R01AG067501, U2C ES030163, R01ES023839, and UL1-TR00187.

## Acknowledgements

We thank Dr. Diane Re for her general feedback throughout the study, and Dr. Vrinda Kalia for general advice regarding *C. elegans* experimentation. Thank you to Dr. Joseph Liang for facilitating the shipping of the UA44 strain to our lab.

## Conflict of interest

The authors have no conflicts of interest to declare.

## Digital Repository

R code used for this study is freely accessible on GitHub through the following link: https://github.com/jdicent1/C.elegans-TCE-Metabolite-Neurotoxicity

All data is accessible through Dryad using the following link: https://doi.org/10.5061/dryad.w6m905r50

## Notes

https://github.com/jdicent1/C.elegans-TCE-Metabolite-Neurotoxicity

https://doi.org/10.5061/dryad.w6m905r50

### Competing Interest Statement

The authors have declared no competing interest.

