## Supplementary Information for "Assessment of dopaminergic neurotoxicity of CYP2E1-mediated trichloroethylene metabolites in *C. elegans*"

### The neurotoxic effects of CYP2E1-mediated trichloroethylene metabolites in *C. elegans*

Affiliations:
[1]Department of Environmental Health Sciences, Mailman School of Public Health at Columbia University, New York, NY, United States.
[2]IARC for Carolina?
[3]Department of Biochemistry and Molecular Biology, Medical University of South Carolina, Charleston, SC, United States
[4]Department of Regenerative Medicine and Cell Biology, Medical University of South Carolina, Charleston, SC, United States
[5]Department of Molecular Pharmacology and Therapeutics, Vagelos College of Physicians and Surgeons, Columbia University, New York, New York, NY, United States.

### SI1: Effect of pH

#### Introduction and Methods

Given the limited knowledge of how pH affects *C. elegans*, we sought to verify whether pH would have a profound effect on experimental results. To avoid confounding experimental results with potential pH effects, we pH-corrected the compounds used in this study to ensure the effects were attributable to the compounds rather than pH. We determined the appropriate solvent and whether pH correction could disentangle pH effects from the effects of the compound itself.

To determine which solvent had the greatest pH-buffering capacity across serial dilutions, we prepared TCA solutions in Milli-Q water, $KPO_4$ buffer, or M9. $KPO_4$ buffer was prepared using 814.088 mg of dibasic potassium phosphate and 44.39 mg of monobasic potassium phosphate in 50 mL of Milli-Q water, yielding a pH of 8. For each solvent, we dissolved the sodium hydroxide used for pH correction in the corresponding solvent. For the highest concentration (6 M), we pH-corrected by adding sodium hydroxide in 1-10 µL increments, then serially diluted to 600 mM, 60 mM, 6 mM, and 3 mM. We verified the pH of all concentrations using pH strips.

For pH effect experiments, we dosed synchronized populations of L1 worms with a single dose of either TCA or pH-adjusted M9 and assessed changes in body size at the L4 stage of development. TCA concentrations ranged from 50 µM to 100 mM, with unadjusted M9 (pH 7.1) as the vehicle control. To isolate the compound's effect from its acidity, we adjusted M9 to pH 5.35, 5.5, 6.0, 6.5, and 7.0 by adding 6 N hydrochloric acid, spanning the pH range of the TCA doses tested. Worms were treated as described above using NGM well plates, and the pH of both the TCA and mock doses was measured with a pH meter.

Worms in NGM-filled 6-well plates were imaged on an APX100 microscope at 4x magnification using brightfield for body size analysis, at consistent light intensity and exposure. Body size was analyzed in ImageJ and Fiji using the segmented line tool to draw a line along the midline of each worm from head to tail. The ROI manager was used to record the lengths of all regions within a single image before exporting. Images were rescaled using an ImageJ macro to convert body length measurements to microns.

Outliers were removed using a ± 2 SD threshold, excluding any worm with a body length more than 2 SDs from the group mean. Data were normalized to the M9 control and expressed as a percentage of the control value before statistical testing. We used Welch's one-way ANOVA, followed by the Games-Howell post hoc test for multiple comparisons. The data analysis was conducted using the following R packages: tidyverse, janitor, readxl, mclust, FactoMineR, factoextra, multcompView, car, afex, rstatix, and DescTools (Wickham and RStudio 2023; Firke et al. 2024; Signorell et al. 2025; Kassambara 2025; Fraley et al. 2025; Singmann et al. 2025; Fox et al. 2026; Graves et al. 2026; Kassambara et al. 2026; Husson et al. 2026; Wickham et al. 2026).

#### Results and Discussion

To determine which solvent would provide the most stable, reproducible pH across serial dilutions, a test of various solvents was performed using TCA (Table SI1.1). Given the highly acidic nature of TCA, the 6 M solution was pH-corrected to a range of 5.5 to 7.5 using 10 M sodium hydroxide. Subsequently, serial dilutions were performed. The pH correction of the TCA led to a significant, non-negligible increase in solution volume. As a result, the 6 M concentration was recalculated to 5.1723 M, the value we used in serial dilution calculations to ensure accurate concentrations.

Milli-Q water had the least buffering capacity, with the pH falling from 7.5 to 6 after 2 dilutions. In comparison, the $KPO_4$-buffered solutions fell from 7.5 to 7 in one dilution but remained at 7 for the remaining dilutions. The M9 solution's starting pH was lower than that of the others, at 5.5, because the equivalence point appears to be around that point in that solvent, as compared to the other solvents. As a result, adding sodium hydroxide beyond this pH raised the pH to 13. Regardless, M9 appears to have a buffering capacity similar to that of the $KPO_4$ buffer, with the pH increasing from 5.5 to 7 after one serial dilution and remaining at 7 for subsequent dilutions. Ultimately, M9 was chosen as the solvent for the remainder of the study because of its buffering capacity and its relevance to worm experiments as a typical laboratory buffer used by worm labs.

| TCA Concentration | Milli-Q Water | $KPO_4$ Buffer | M9 |
|---|---|---|---|
| 5.1723 M | 7.5 | 7.5 | 5.5 |
| 600 mM | 7 | 7 | 7 |
| 60 mM | 6 | 7 | 7 |
| 6 mM | 6 | 7 | 7 |
| 3 mM | 6 | 7 | 7 |

**Table SI1.1.** pH values of serially diluted TCA solutions using Milli-Q water, $KPO_4$, or M9 as the solvent.

After selecting M9 as the solvent, we scaled up dose volumes and concentrations for dosing experiments. Table SI1.2 shows the final pH for each TCA concentration. We then used this pH range to prepare mock M9 solutions to assess whether pH alone affected worm body size, independent of the compound. We used select TCA doses to validate the dosing approach and test the dosing medium.

| Initial TCA Concentration | pH of Initial TCA Concentration | TCA Concentration on Well Plate |
|---|---|---|
| 5.1723 M | 5.35 | 86.205 mM |
| 3 M | 6.09 | 50 mM |
| 1.8 M | 6.49 | 30 mM |
| 600 mM | 6.78 | 10 mM |
| 300 mM | 6.92 | 5 mM |
| 180 mM | 6.99 | 3 mM |
| 60 mM | 7.05 | 1 mM |
| 30 mM | 7.08 | 500 µM |
| 18 mM | 7.12 | 300 µM |
| 6 mM | 7.12 | 100 µM |
| 3 mM | 7.09 | 50 µM |
| M9 | 7.09 | N/A |

**Table SI1.2.** pH values of serially diluted TCA solutions in M9 and the concentrations they were used to create on the well plates.

To determine whether pH alone significantly affected worm body size, mock doses were prepared using M9 acidified with hydrochloric acid to pH values within the range observed across the TCA concentrations in Table SI1.2 (Figure SI1.1). Of the pH levels tested, only pH 6.5 produced a statistically significant difference from the M9 control (pH 7.1); worms treated at pH 6.5 were, on average, larger than those in the M9 control (Table SI1.3).

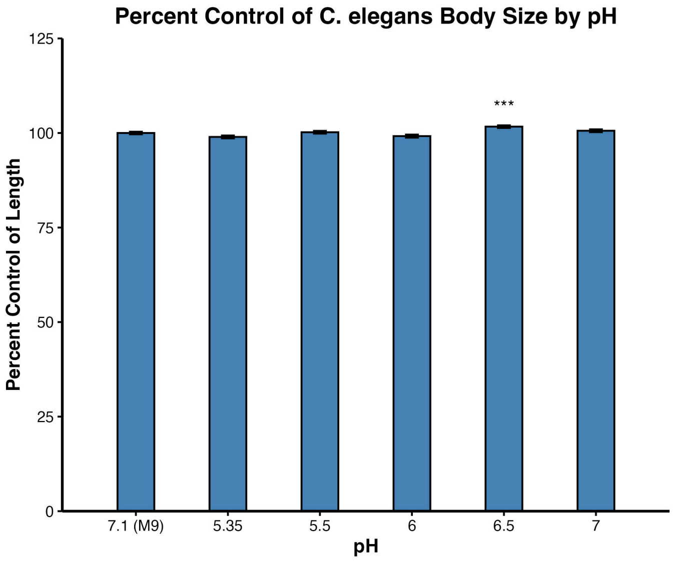


**Figure SI1.1.** Mean percent control of *C. elegans* body size following exposure to M9 buffer at varying pH levels. Dosing was conducted using 6-well plates. Body size was measured using manual analysis in ImageJ and Fiji. Error bars represent ± SEM. Levene's test indicated heteroscedasticity ($p < 0.05$), so Welch's one-way ANOVA was used ($W(5, 1338.2) = 10.749$, $p < 0.0001$). A significant pairwise difference was observed between pH 6.5 and the M9 control (pH 7.1) (Games-Howell, $p < 0.001$). Asterisks indicate significant difference from the M9 control (Games-Howell; *$p < 0.05$, **$p < 0.01$, ***$p < 0.001$). n = 432 - 523 worms per treatment, across 3 experimental replicates.

| Contrast | Difference | Lower 95% CI | Upper 95% CI | p-value |
|---|---|---|---|---|
| 5.35 - M9 | -1.04 | -2.24 | 0.16 | 0.13 |
| 5.5 - M9 | 0.20 | -0.96 | 1.36 | 1.00 |
| 6 - M9 | -0.83 | -2.08 | 0.43 | 0.42 |
| 6.5 - M9 | 1.68 | 0.55 | 2.81 | < 0.001 |
| 7 - M9 | 0.60 | -0.59 | 1.78 | 0.71 |

**Table SI1.3.** Games-Howell post-hoc test results comparing the percent control of worm body size across pH levels to the M9 control. 95% confidence intervals and family-wise adjusted p-values are reported.

As a proof of concept, we exposed worms to pH-corrected doses of the compound to determine whether changes in body size were attributable to the compound rather than to pH (Figure SI1.2). Body size was significantly altered following TCA exposure compared to the M9 control, despite no observed difference attributable to pH at comparable pH levels. Body size decreased significantly relative to the control at final well plate concentrations of 3 mM or higher (Table SI1.4). Notably, although pH 6.5 falls within the pH range corresponding to the 3 mM to 86.205 mM doses (pH 6.99 to 5.35), the mock pH experiments showed the opposite effect of an increase rather than a decrease in body size. This suggests that the decreases observed at these concentrations are attributable to TCA rather than pH. A significant increase in body size relative to the control was also observed at 300 µM and 500 µM, both with neutral pH values similar to the control, further suggesting that the increases at these concentrations are attributable to TCA rather than pH.

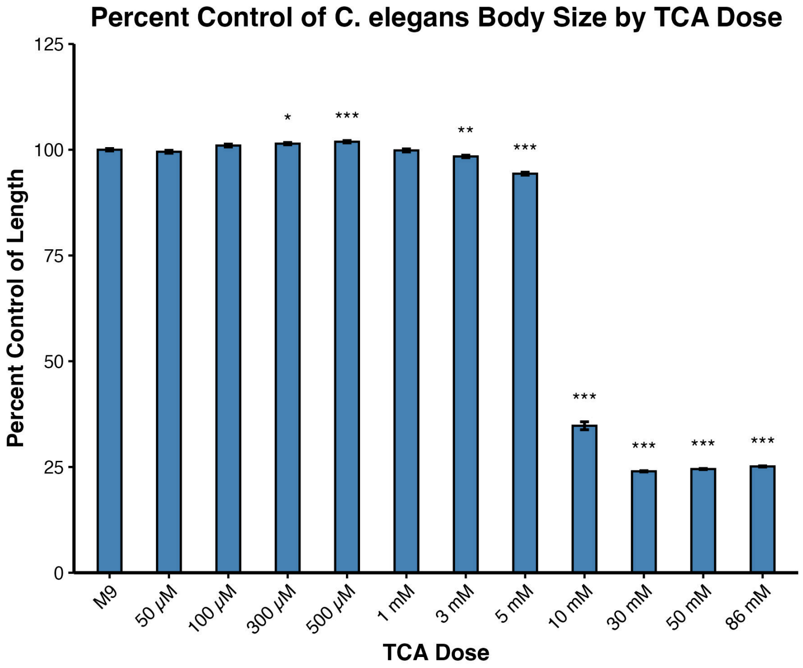


**Figure SI1.2.** Mean percent control of *C. elegans* body size following exposure to TCA at varying concentrations. Dosing was conducted using 6-well plates. Body size was measured using manual analysis in ImageJ and Fiji. Error bars represent ± SEM. Welch's one-way ANOVA was used ($W(11, 1895.7) = 28730$, $p < 0.0001$). Asterisks indicate significant difference from the M9 control (Games-Howell; *$p < 0.05$, **$p < 0.01$, ***$p < 0.001$). A significant pairwise difference was observed between 300 µM, 500 µM, 3 mM, 5 mM, 10 mM, 30 mM, 50 mM, and 100 mM doses compared to the M9 control. (Games-Howell, $p < 0.001$). n = 255 - 504 worms per treatment, across 3 experimental replicates.

| Contrast | Difference | Lower 95% CI | Upper 95% CI | p-value |
|---|---|---|---|---|
| 50 µM - M9 | -0.475 | -1.92 | 0.972 | 0.996 |
| 100 µM - M9 | 0.997 | 0.449 | 2.44 | 0.507 |
| 300 µM - M9 | 1.443 | 0.165 | 2.72 | **0.012** |
| 500 µM - M9 | 1.896 | 0.602 | 3.19 | **< 0.001** |
| 1 mM - M9 | -0.157 | -1.65 | 1.34 | 1.0000 |
| 3 mM - M9 | -1.582 | -2.92 | -0.243 | **0.006** |
| 5 mM - M9 | -5.657 | -7.12 | -4.20 | **< 0.001** |
| 10 mM - M9 | -65.254 | -68.5 | -62.0 | **< 0.001** |
| 30 mM - M9 | -76.022 | -77.1 | -75.0 | **< 0.001** |
| 50 mM - M9 | -75.487 | -76.5 | -74.5 | **< 0.001** |
| 86 mM - M9 | -74.868 | -75.9 | -73.8 | **< 0.001** |

**Table SI1.4.** Games-Howell post-hoc test results comparing the percent control of worm body size across TCA doses to the M9 control. 95% confidence intervals and family-wise adjusted p-values are reported.

M9 was selected as the best buffer pH among those tested. Pure water has no buffering capacity because it has no salts of weak conjugate acids or bases to neutralize compounds. The $KPO_4$ buffer contained a mixture of monobasic potassium phosphate ($KH_2PO_4$) and dibasic potassium phosphate ($K_2HPO_4$), resulting in a pH of 8. We chose a $KPO_4$ buffer because phosphates have a high buffering capacity. Because the buffer started at the highest pH among the tested solvents, with M9 and water both at pH 7, it was reasonable to expect a relatively neutral pH at the highest concentration and good buffering throughout the serial dilutions. M9, on the other hand, contains monobasic potassium phosphate ($KH_2PO_4$), sodium phosphate ($Na_2HPO_4$), sodium chloride (NaCl), and magnesium sulfate ($MgSO_4$). Because it contains phosphates, it has a buffering capacity similar to that of the $KPO_4$ buffer. Sulfate, as the conjugate base of a strong acid (sulfuric acid or $H_2SO_4$), has practically no buffering capacity since the sulfate ion completely dissociates from the hydrogen ions in solution. Regardless, M9 is commonly used in labs working with *C. elegans*, giving us another reason to select M9 over $KPO_4$ buffer as our solvent.

After deriving a range of pH-corrected TCA values, we found that pH values within this range did not substantially affect body size. The only exception was pH 6.5, at which we observed a slight increase in body size compared with worms dosed with the M9 control. Increased body size in response to an exposure is uncommon; typically, we see that worms increase body size in response to genetic mutations that affect key regulators of body size. For example, studies have found that *che-2*, *osm-6*, and *che-3* mutants, which have structural defects in cilium structure, have altered sensory perception and regulation of growth to a normal body size; *egl-4* mutants, which fail to repress transforming growth factor-β (TGF-β) signaling, experience overactivity of the TGF-β pathway leading to overgrowth (Fujiwara et al. 2002; Hirose et al. 2003). *cat-2* mutants, which have defects in the rate-limiting enzyme for dopamine synthesis, tyrosine hydroxylase, have decreased dopamine levels and an increased body length and width (Nagashima et al. 2016). Typical studies that expose C. elegans to an environmental exposure of interest, such as ethanol, bisphenol A (BPA), or pesticides, tend to show decreased body size (Davis et al. 2008; Ruan et al. 2009; Zhou et al. 2016). Therefore, we found a slight increase. One possible explanation is that a pH of 6.5 may affect the worms' feeding behavior or motility, leading to increased body size. Another possible explanation is that our large sample size (432 - 523 worms per treatment), which is substantially greater than most worm studies that do not use a COPAS, gives us greater power to detect small, statistically significant differences. In any case, we ultimately continued to use remade, pH-corrected doses, with pH values all around 7.

Despite an increase in body size at pH 6.5, pH-corrected TCA still decreased worm body size at concentrations near 6.5 (10 and 30 mM). This is consistent with the general pattern of toxicant-induced size decreases in *C. elegans*. We also observed a small but statistically significant increase in body size at 300 and 500 µM, with percent differences relative to the control of 1.443% and 1.896%, respectively. Again, this raises the possibility that the large sample size of 255-504 worms per treatment provides sufficient statistical power to detect small differences. Regardless, this experiment provides conceptual proof that when doses are pH-corrected to a more neutral range, the effects on body size are more likely attributable to the compound than to pH.

### SI2: Dosing Medium

#### Introduction and Methods

Since our laboratory had not previously used NGM-filled well plates for *C. elegans* experiments, a series of tests was conducted to determine their suitability. First, it was determined whether the volume of NGM in the wells affected the exposure. 3 mL of agar was the minimum required to comfortably fill the bottom of the well, so volumes from 3 mL to 8 mL in 1 mL increments were tested. Then, we tested the difference between using liquid culture, NGM-filled well plates, or 6-cm NGM plates for dosing using doses of TCA known to cause an effect. Finally, we developed a filtering method for more precise data measurement while using the Complex Object Parametric Analyzer and Sorter (COPAS), which was used to streamline these analyses.

L1 worms were dosed with a single dose of TCA and assessed at the L4 stage of development for changes in body size. TCA concentrations ranged from 1 mM to 100 mM, with M9 as the vehicle control. For NGM-filled 6-well plate and 6 cm plate dosing, the treatment solution was pipetted directly onto the agar in small droplets and allowed to equilibrate for 30 minutes before worms were added. For liquid culture, the treatment was mixed with food, S-complete medium, and ampicillin, then distributed to the wells of a 24-well plate.

Worms were analyzed using the Complex Object Particle Analyzer and Sorter Flowpilot-250 (COPAS FP-250), which was purchased from Union Biometrica (Holliston, MA). Worms were washed from plates using Milli-Q water and transferred to 1.5 mL microcentrifuge tubes. Worms that were in liquid culture were transferred to 1.5 mL microcentrifuge tubes. Worms were allowed to settle at the bottom of the tubes and were washed twice with Milli-Q water. The final suspension of worms was in water with 5 mM levamisole to anesthetize them prior to analysis by the COPAS. For length analysis using the N2 strain, time-of-flight (TOF), which represents worm length, was used as the primary outcome measure. These files were exported to a .csv file and analyzed using RStudio.

A filtering method was developed to screen out false positives in the COPAS data. For example, some treated worms began laying eggs before the end of the experiment, resulting in L1s and unhatched eggs being measured alongside the treated generation. Since the population of interest was the treated generation, it was important to exclude these smaller objects. This was particularly challenging because some treatments caused severe decreases in body size, such that developmentally delayed worms at the L4 stage were occasionally comparable in size to L1s. To address this, a filtering pipeline was developed to remove eggs and L1s across all treatments without substantially reducing the number of worms retained in treatments with severe developmental defects.

To maintain an objective approach, data were split by experiment. Within each experiment, principal components analysis (PCA) and k-means clustering with three clusters were applied to group the data. The PCA was conducted using extinction, TOF, peak width of extinction, and peak width of TOF. Clusters were then visually assessed, and those with COPAS parameters of extinction (representing worm width) and TOF closest to zero were excluded. The filtering pipeline is illustrated in Figure SI2.1. Briefly, data points within each experiment were plotted following PCA-based k-means clustering (Figure SI2.1A), the cluster closest to the origin was removed (Figure SI2.1B), and the remaining points were faceted by dose or treatment for a final visual inspection of the worm population across conditions (Figure SI2.1C).

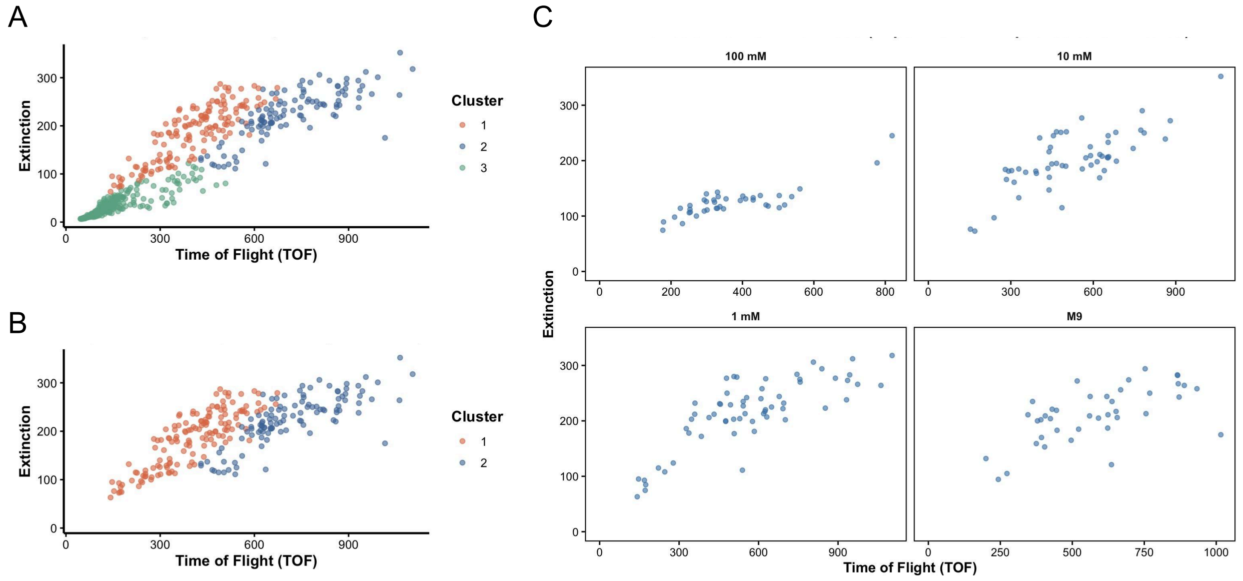


**Figure SI2.1.** Representative plots illustrating the COPAS data filtering pipeline. (A) Extinction vs. TOF graph for experiment 1 of the liquid culture replicates, prior to filtering. Points are colored by cluster as determined by the PCA-based k-means clustering (k = 3). (B) Extinction vs. TOF graph for experiment 1 of the liquid culture replicates, following the removal of cluster with values closest to the origin (Cluster 3). (C) Extinction vs. TOF graph for experiment 1 of the liquid culture replicates, post-filtering, faceted by dose.

Data were normalized to the M9 control and expressed as a percentage of the control value before statistical testing. Normalization was performed within each experiment and dosing medium to account for inter-experiment variability. Two-way ANOVA with White's correction for heteroscedasticity or Welch's one-way ANOVA was used. The Games-Howell post hoc test was then used for multiple comparisons. Coefficients of variation were also assessed for the COPAS filtering pipeline. All data filtering and statistical analysis were conducted in RStudio (Version 2025.09.2+418) using the following packages: tidyverse, janitor, readxl, mclust, FactoMineR, factoextra, multcompView, car, afex, rstatix, and DescTools (Wickham and RStudio 2023; Firke et al. 2024; Signorell et al. 2025; Kassambara 2025; Fraley et al. 2025; Singmann et al. 2025; Fox et al. 2026; Graves et al. 2026; Kassambara et al. 2026; Husson et al. 2026; Wickham et al. 2026).

#### Results and Discussion

Since our laboratory had not previously used NGM-filled well plates for *C. elegans* experiments, a series of tests was conducted to determine their suitability. First, it was determined whether the volume of NGM in the wells affected the exposure (Figure SI2.2). Prior to using the filtering method developed for the COPAS data, there were significant differences in agar volumes for all volumes except 3 mL (Figure SI2.2A). After the COPAS filtering pipeline, which minimized the number of significant differences between agar volumes from 5 to 2, 3 mL and 5 mL had no significant differences with other agar volumes (Figure SI2.2B). Since the 3 mL volume was the only one that was not significantly different from any other agar volume pre- and post-filtering, it was selected for use for all future experiments. In addition, the 3 mL volume was the only volume in which worms were within the focal plane of the objective without removing them from the well plate. The use of NGM-filled, low-evaporation well plates provided a high-throughput option for dosing for assays that require the worms to be on NGM, as well as for imaging.

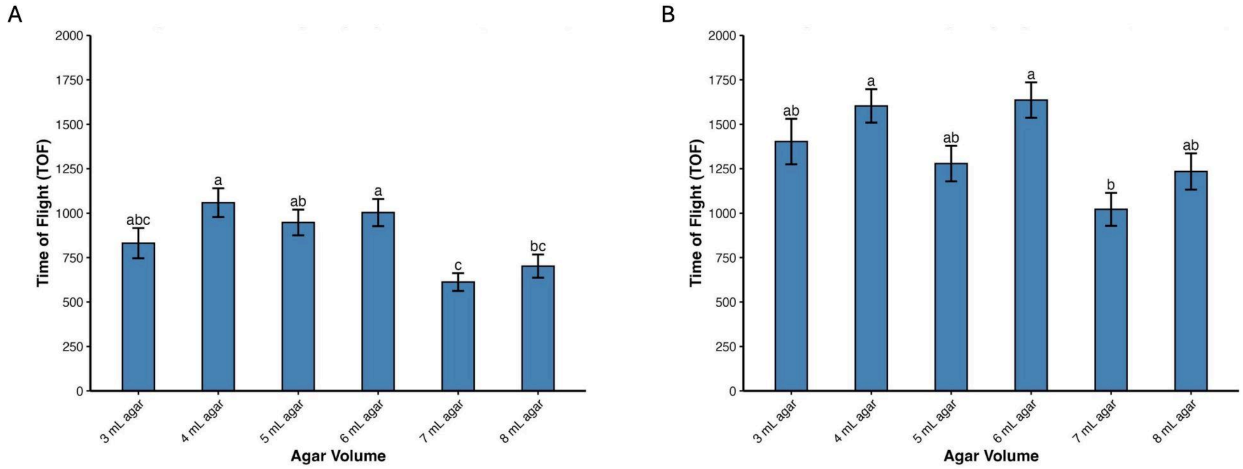


**Figure SI2.2.** Mean time of flight (TOF) of *C. elegans* treated with 10 mM TCA on varying agar volumes in a 6-well plate, before and after filtering. Error bars represent ± SEM. Bars that share a letter are not significantly different from one another (Games-Howell, $p < 0.05$). (A) Pre-filtered means of TOF of *C. elegans*. Welch's one-way ANOVA ($W(5, 390.61) = 7.54$, $p < 0.001$). Significant pairwise differences were observed for 4 mL vs 7 mL, 4 mL vs. 8 mL, 5 mL vs. 7 mL, 6 mL vs. 7 mL, and 6 mL vs. 8 mL (Games-Howell, $p < 0.05$). n = 106 - 202 worms per agar volume, across 2 experimental replicates. (B) Post-filtered means of TOF of *C. elegans*. Welch's one-way ANOVA ($W(5, 205.79) = 5.937$, $p < 0.001$). Significant pairwise differences were observed for 4 mL vs. 7 mL and 6 mL vs. 7 mL (Games-Howell, $p < 0.05$). n = 52 - 94 worms per agar volume, across 2 experimental replicates.

To determine whether dosing format influences worm response, three TCA concentrations (100 mM, 10 mM, and 1 mM) were tested across 6 cm plate, NGM-filled 6-well plate, and liquid culture formats using the COPAS for high-throughput data acquisition (Figure SI2.3). Since TOF values were normalized to each format's own M9 control, these comparisons reflect differences in dose-response patterns rather than absolute body size. Significant pairwise differences were observed across all three formats for worms treated with 100 mM TCA. Significant pairwise differences were also observed between the 6 cm plate and liquid culture and between the 6-well plate and liquid culture for worms treated with 10 mM TCA. At the 1 mM dose, a significant difference was observed between the 6-cm plate and 6-well plate conditions. Liquid culture produced the smallest reduction in body size at higher doses, whereas 6-cm plate dosing generally produced the greatest reduction relative to the control.

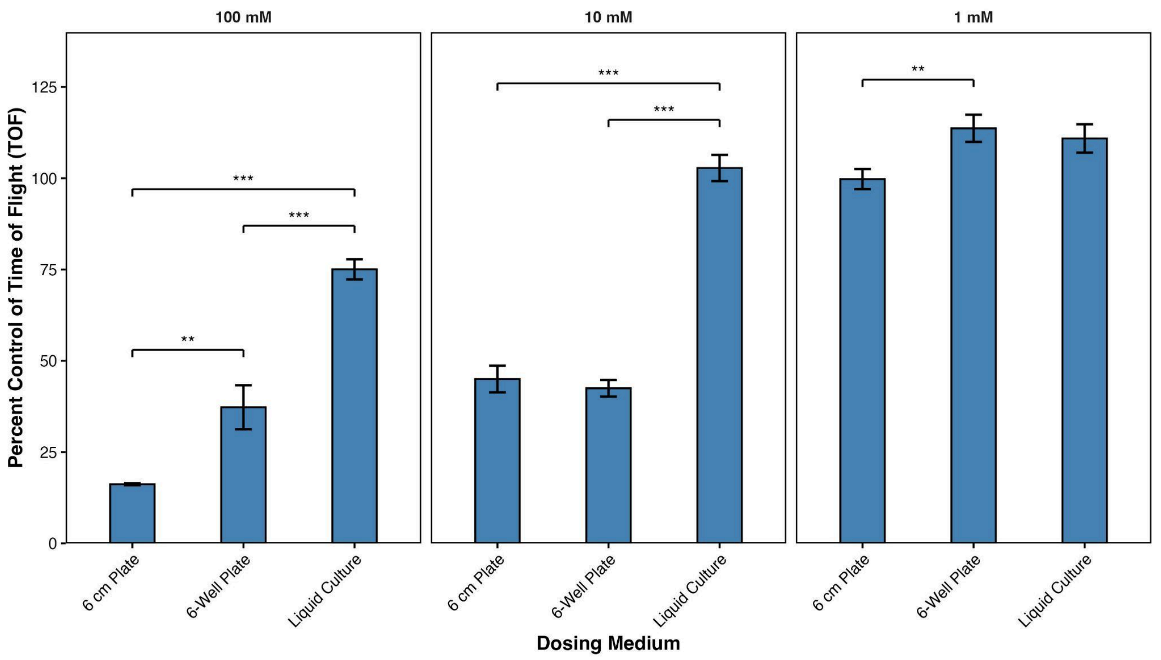


**Figure SI2.3.** Mean percent control of TOF of *C. elegans* dosed with 100 mM, 10 mM, or 1 mM TCA across dosing media. Percent control values were calculated by normalizing individual worm TOF values to the mean TOF of the M9 control within each medium and experiment. Error bars represent ± SEM. Two-way ANOVA with White's correction for heteroscedasticity was used: dose ($F(3, 3511) = 152.12$, $p < 0.001$), medium ($F(2, 3511) = 56.63$, $p < 0.001$), dose × medium interaction ($F(6, 3511) = 21.40$, $p < 0.001$). Asterisks indicate significant pairwise differences between media within each dose (Games-Howell; *$p < 0.05$, **$p < 0.01$, ***$p < 0.001$). Significant pairwise differences were observed across all media for worms treated with the 100 mM TCA, across the 6 cm plate vs. liquid culture conditions and 6-well plate vs. liquid culture conditions for the worms treated with 10 mM TCA, and across the 6 cm plate vs. 6-well plate conditions for the worms treated with 1 mM TCA (Games-Howell, $p < 0.05$). M9 control values did not differ significantly across dosing media (Games-Howell, $p = 1.00$ for all comparisons) and are not shown. n = 99 - 597 worms per dose per medium across 2 experimental replicates.

To determine whether the COPAS filtering pipeline appropriately filtered the dataset, the coefficient of variation (CV) and the number of worms retained per dose was examined. The CV was assessed as a measure of data spread; the assumption was that if the CV decreased post-filtering, the COPAS filtering pipeline successfully reduced the spread of values and removed a significant portion of eggs and L1s. In general, the pipeline decreased the CV across all dosing media and agar volumes tested (Tables SI2.1 and SI2.2). The exceptions to this were the worms dosed with 100 mM TCA in both the 6 cm plate and 6-well plate formats, where the

CV increased after the filtering pipeline. The CV was generally lower for the liquid culture dosing medium than for the other dosing media. The 6 cm plate and 6-well plate dosing media had relatively similar CV values pre- and post-filtering. In addition, CV values for the different agar volumes within a 6-well plate were relatively similar pre- and post-filtering.

This filtering pipeline reduced the number of worms included across all doses. The 100 mM TCA dose showed a more pronounced reduction than the other doses, regardless of the number of experimental replicates. In general, the filtering pipeline was able to exclude eggs and L1s from the experiments without completely excluding all worms at the highest doses across most conditions. There was one instance in which the 100 mM dose was completely excluded by the filtering pipeline, corresponding to Experiment 2 of the 6-well plate replicates.

| Dosing Medium | Dose | N Pre-Filtering | N Post-Filtering | CV Pre-Filtering (%) | CV Post-Filtering (%) | CV Difference (%) |
|---|---|---|---|---|---|---|
| 6 cm Plate | 100 mM | 314 | 126 | 35.79 | 18.49 | -17.30 |
| | 10 mM | 353 | 152 | 100.07 | 100.03 | -0.04 |
| | 1 mM | 1110 | 597 | 117.00 | 67.24 | -49.76 |
| | M9 | 1118 | 599 | 116.25 | 65.31 | -50.94 |
| 6-Well Plate | 100 mM | 328 | 99 | 152.57 | 161.12 | 8.55 |
| | 10 mM | 1431 | 510 | 140.19 | 121.20 | -18.98 |
| | 1 mM | 652 | 326 | 118.92 | 59.26 | -59.66 |
| | M9 | 818 | 382 | 131.54 | 68.21 | -63.33 |
| Liquid Culture | 100 mM | 221 | 105 | 62.51 | 37.49 | -25.02 |
| | 10 mM | 422 | 213 | 79.75 | 51.20 | -28.55 |
| | 1 mM | 686 | 258 | 85.98 | 56.43 | -29.55 |
| | M9 | 391 | 156 | 77.59 | 43.88 | -33.71 |

**Table SI2.1.** Coefficient of variation (CV) of percent control values for each dose within each dosing medium before and after filtering. The difference is reported as the post-filtering CV minus the pre-filtering CV, where a negative value indicates reduced variability after filtering. N for the 6 cm plate and liquid culture is compiled from 2 experimental replicates; N for the 6-well plates is compiled from 4 experimental replicates.

| Volume of Agar (mL) | N Pre-Filtering | N Post-Filtering | CV Pre-Filtering (%) | CV Post-Filtering (%) | CV Difference (%) |
|---|---|---|---|---|---|
| 3 | 106 | 52 | 103.18 | 64.16 | -39.02 |
| 4 | 131 | 77 | 88.89 | 53.27 | -35.62 |
| 5 | 140 | 86 | 91.41 | 75.90 | -15.51 |
| 6 | 174 | 94 | 98.57 | 57.93 | -40.64 |
| 7 | 202 | 89 | 118.07 | 89.31 | -28.76 |
| 8 | 154 | 74 | 114.64 | 76.47 | -38.18 |

**Table SI2.2.** Coefficient of variation (CV) of percent control values for the test of each potential agar volume in a 6-well plate before and after filtering. The difference is reported as the post-filtering CV minus the pre-filtering CV, where a negative value indicates reduced variability after filtering. N is compiled from 2 experimental replicates.

The *C. elegans* response to a single 10 mM TCA dose varied with agar volume. Using 3.5 mL of agar is common in studies that use 6-well plates with *C. elegans*; however, few studies use 6-well plates, and there is no experimentally validated field-wide gold standard for this volume. Thus, it was reasonable to select the agar volume that was not significantly different from the others, 3 mL, as the best option. The selection of 3 mL was also a logistical choice, as it allowed the worms to remain in the focal plane for imaging in downstream microscopy experiments. Any volume above 3 mL resulted in difficulty focusing on the worms or an inability to focus on them entirely.

There is limited literature on the observed differences between dosing with agar plates as opposed to liquid culture, though it is referenced throughout the literature that liquid culture provides greater control over experimental variables, food, and dosing (Win et al. 2013; Pittman et al. 2017; Hibshman et al. 2021; Wang et al. 2023). These results were consistent with the literature, as the coefficient of variation for the liquid culture experiments was much lower than those for the 6-well plates or the agar plates, indicating that dosing was more uniform across worms in liquid culture. Worms in liquid culture also appeared more resilient to changes in body size at high doses than those in other dosing formats. This could be due to variations in dose uniformity: in the agar-based dosing formats, compound absorption into the agar may be variable. This could result in some worms receiving disproportionately high doses, increasing variability and driving down the group means.

Despite its advantages, liquid culture is not always the most appropriate dosing format. For example, *C. elegans* release all their eggs into solution in liquid culture, which can confound or render impossible any reproduction assay. In this instance, it would be more appropriate to use agar plates or 6-well plates, even if they may not provide as uniform dosing as liquid culture. Notably, worms are more resilient to change in 6-well plates than on agar plates. This may reflect that the smaller amount of agar per well on the 6-well plates results in a more uniform distribution of the compound than in the 6 cm plates. Regardless, 6-well plates provide convenience and efficiency for many assays, making them a better option than agar plates in many experimental contexts.

A reliable pipeline for filtering COPAS data was successfully developed, though there are several caveats. Although the pipeline is designed to be objective by applying the same filtering criteria across all doses within an experiment, it tends to exclude a greater proportion of worms that have received a dose that causes severe developmental changes, because these worms can be similar in size to eggs or L1s. Notably, when the treatment is uniform across an experiment, as in the agar volume experiment, a similar proportion of worms was excluded across replicates. This suggests that the pipeline performs consistently under these conditions. In dose-response experiments, however, it is more likely that the pipeline excludes a variable number of worms across doses with varying effects. The number of worms retained after filtering may be worth noting. A dose that retains very few worms post-filtering may indicate that few worms reached a developmentally appropriate size and can inform the severity of the effect, proportional to the number of worms initially introduced into the filtering pipeline.

The pipeline also offers flexibility in the number of clusters used in k-means, which can be tailored to the experimental context. Overall, reductions in coefficient of variation following filtering suggest that the pipeline improves data uniformity while preserving the underlying biological signal. The COPAS approach is also substantially higher-throughput than microscopy-based body size assessment, though the latter provides greater precision.

### SI3: Lethality Assessments

#### Introduction and Methods

We conducted a lethality assay to confirm the appropriate single doses for the main study. We conducted two separate studies. In the first study, we dosed N2, BZ555, or UA44 worms with vehicle control (M9), solvent control (M9 with 0.3% DMSO), TCA, or TCEOH. The tested TCA concentrations were 100 mM, 50 mM, 25 mM, 10 mM, 5 mM, 2.5 mM, and 1 mM. Because of restrictions on TCEOH dissolution, we did not test 100 mM.
Using liquid culture and a 48-well plate, we dosed L4 worms with the desired treatment and assessed compound lethality on day 1 and day 3 of adulthood, refreshing the dose on day 2 of adulthood. We used separate wells for the worms assessed each day.

Each day, we manually counted live and dead worms. Using Google Sheets, we calculated the percentage of live worms. We graphed these percentages in GraphPad Prism and conducted two-way ANOVAs with Tukey's post hoc test.

#### Results and Discussion

We observed that the highest doses yielded the lowest percentages of live worms. For TCA, dead worms were observed on days 1 and 3 at 100 mM and 50 mM concentrations across all worm strains (Figure SI3.1). UA44 worms appeared particularly sensitive to these doses compared with the other strains. We observed differences between the 25 mM and 10 mM doses in the percentage of live worms, and we aimed to select the highest dose with minimal to no lethality. We ultimately chose 10 mM because it was slightly more consistent across worm strains than 25 mM.

For TCEOH, dead worms were observed on days 1 and 3 at 10 mM, 25 mM, and 50 mM concentrations across all worms (Figure SI3.2). We observed a clear difference in the percentage of live worms between the 10 mM and 5 mM doses, so we ultimately chose 5 mM for our experiments. Although the 5 mM dose had some dead worms, it was no different from the control doses.

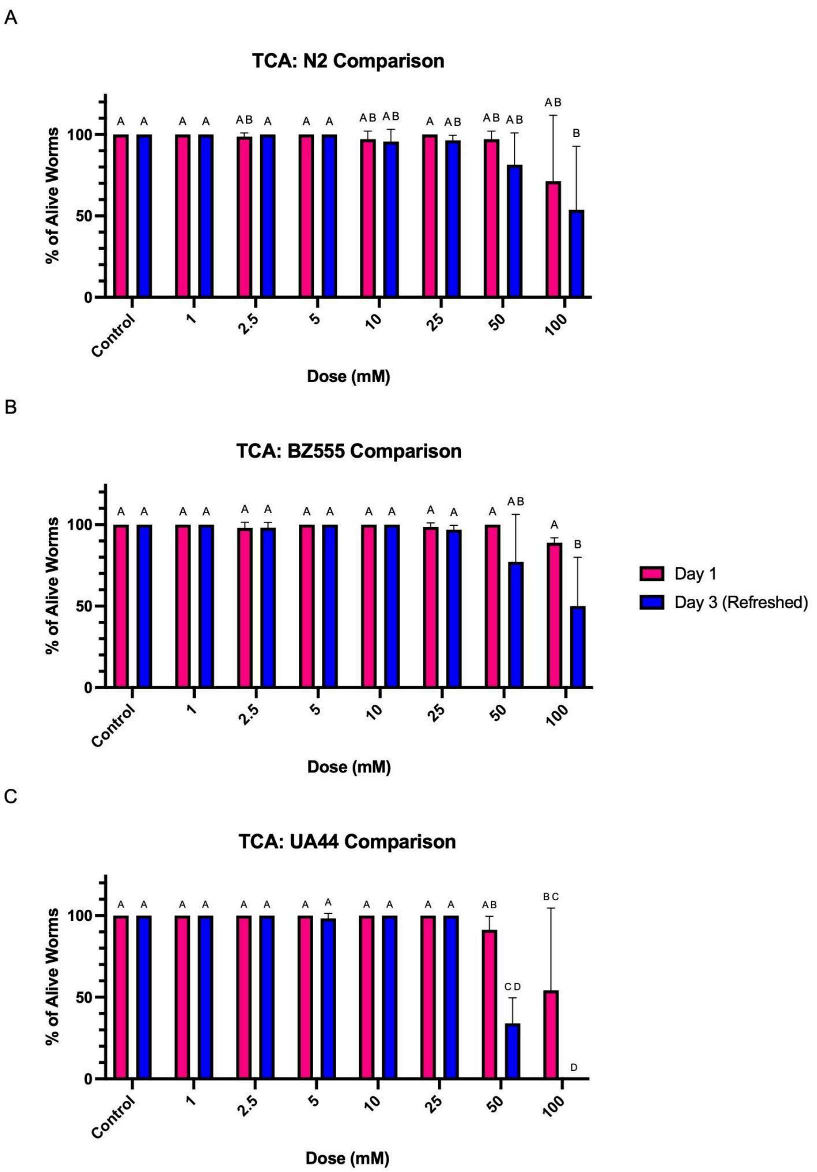


**Figure SI3.1.** Percent of alive worms on day 1 and day 3 of adulthood, with refreshing the dose at day 2 of adulthood, in (A) N2, (B) BZ555, and (C) UA44 worms using TCA. Two-way ANOVA was used, followed by Tukey's post-hoc test for multiple comparisons. Error bars represent ± SD. Bars that share a letter are not significantly different from one another (Tukey's, $p < 0.05$). N2: two-way ANOVA revealed a significant effect of Dose ($F(7, 32) = 4.370$, $p = 0.0017$), but no significant effect of Age ($F(1, 32) = 1.117$, $p = 0.2984$) and no significant Dose × Age interaction ($F(7, 32) = 0.376$, $p = 0.9097$). BZ555: two-way ANOVA revealed significant effects of Dose ($F(7, 32) = 6.151$, $p = 0.0001$) and Age ($F(1, 32) = 6.699$, $p = 0.0144$), as well as a significant Dose × Age interaction ($F(7, 32) = 2.940$, $p = 0.0170$). UA44: two-way ANOVA revealed significant effects of Dose ($F(7, 32) = 24.64$, $p < 0.0001$) and Age ($F(1, 32) = 13.37$, $p = 0.0009$), as well as a significant Dose × Age interaction ($F(7, 32) = 5.505$, $p = 0.0003$). n = 54 - 88 worms per dose, across three experimental replicates.

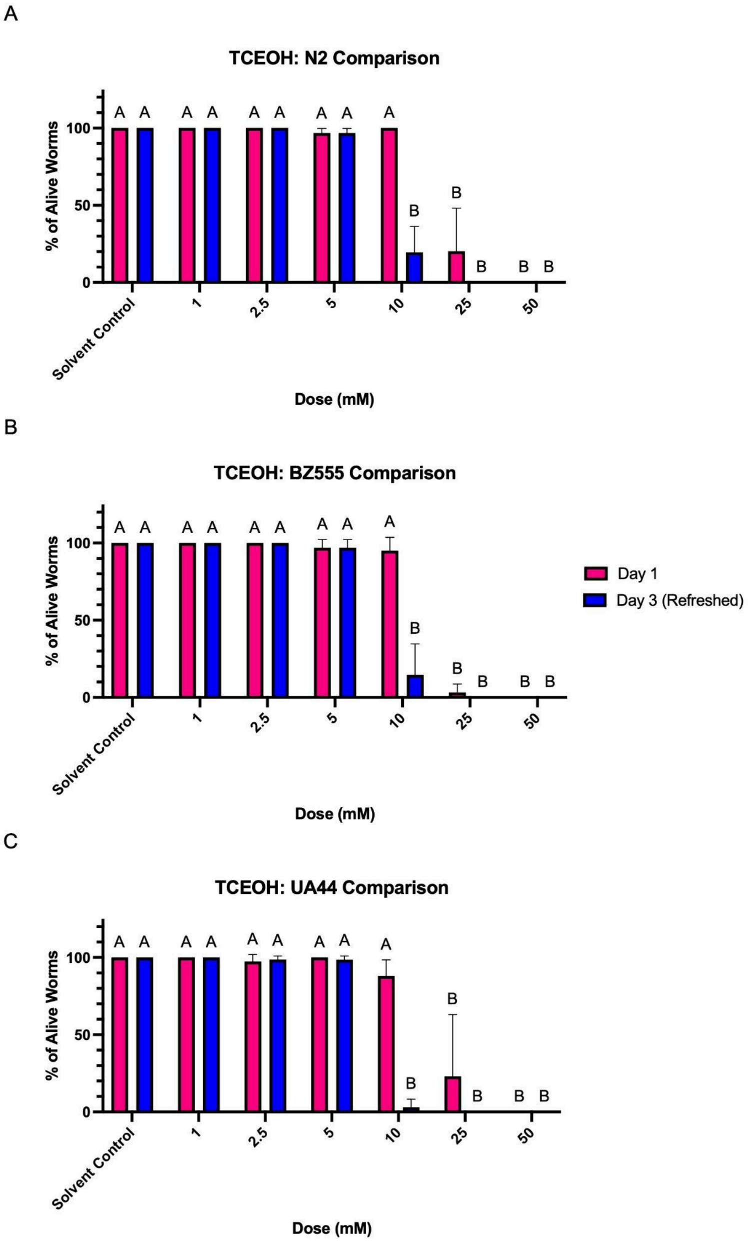


**Figure SI3.2.** Percent of alive worms on day 1 and day 3 of adulthood, with refreshing the dose at day 2 of adulthood, in (A) N2, (B) BZ555, and (C) UA44 worms using TCEOH. Two-way ANOVA was used, followed by Tukey's post-hoc test for multiple comparisons. Error bars represent ± SD. Bars that share a letter are not significantly different from one another (Tukey's, $p < 0.05$). N2: two-way ANOVA revealed significant effects of Dose ($F(6, 28) = 154.8$, $p < 0.0001$) and Age ($F(1, 28) = 28.19$, $p < 0.0001$), as well as a significant Dose × Age interaction ($F(6, 28) = 17.68$, $p < 0.0001$). BZ555: two-way ANOVA revealed significant effects of Dose ($F(6, 28) = 323.9$, $p < 0.0001$) and Age ($F(1, 28) = 37.17$, $p < 0.0001$), as well as a significant Dose × Age interaction ($F(6, 28) = 34.00$, $p < 0.0001$). UA44: two-way ANOVA revealed significant effects of Dose ($F(6, 28) = 96.72$, $p < 0.0001$) and Age ($F(1, 28) = 19.94$, $p = 0.0001$), as well as a significant Dose × Age interaction ($F(6, 28) = 12.10$, $p < 0.0001$). $n = 51$ - $101$ worms per dose, across three experimental replicates.

Taken together, these results enabled us to select appropriate doses for the experiments presented in the main text. For TCA, we had to decide which dose to use, so we carefully reviewed the error bars and consistency. For TCEOH, the decision was simpler, as we noticed a marked difference between the two doses and selected the dose closest to the dose where we see lethality.

### SI4: ImageJ Macros

Below is the code used for the 4x, 20x, and 60x objective images in the ImageJ Macros. These macros were generated using ChatGPT (ChatGPT), due to unfamiliarity with the ImageJ Macro (IJM) language, which is Java-based. The macro code for 4x images was used for the length analyses; the macro code for the 20x and 60x images was used for the neuronal cell body and the dendrite morphology analyses, respectively.

**4x Images - ImageJ Macro:**

```
// Dialog to choose input and output folders
inputDir = getDirectory("Choose Input Directory");
outputDir = getDirectory("Choose Output Directory");

// Get list of all files in the input directory
list = getFileList(inputDir);

// Process each file in the input directory
for (i = 0; i < list.length; i++) {
   // Ignore directories and non-image files
   if (endsWith(list[i], ".tif") || endsWith(list[i], ".tiff")) {
      // Debug: Print current file being processed
      print("Processing: " + list[i]);

      // Open the image
      open(inputDir + list[i]);

      // Select the image (optional as open will do this)
      selectWindow(list[i]);

      // Set the scale
      run("Set Scale...", "distance=0.6154 known=1 unit=microns");

      // Modify the output file name to include '_rescaled'
      fileNameWithoutExtension = list[i].substring(0, list[i].lastIndexOf("."));
      newFileName = fileNameWithoutExtension + "_rescaled.tif";

      // Save the processed image in the output directory
      saveAs("Tiff", outputDir + newFileName);

      // Close the image
      close();
```

```
        // Debug: Print successful processing message
        print("Successfully processed: " + list[i] + " as " + newFileName);
    } else {
        // Debug: Print skipping message for non-image files
        print("Skipping non-image file: " + list[i]);
    }
}
```

**20x Images - ImageJ Macro:**

```
// Ask for folders
inputDir = getDirectory("Input_folder");
if (inputDir == "") exit("No input folder selected.");

outputDir = getDirectory("Output_folder");
if (outputDir == "") exit("No output folder selected.");

// Create subfolders for MIP and Binary if they don't exist
mipDir = outputDir + "MIP/";
binaryDir = outputDir + "Binary/";
File.makeDirectory(mipDir);
File.makeDirectory(binaryDir);

// Get file list
list = getFileList(inputDir);

setBatchMode(true);

for (i = 0; i < list.length; i++) {

    fileName = list[i];

    // Only .tif/.tiff files containing "20x"
    if ((endsWith(fileName, ".tif") || endsWith(fileName, ".tiff")) &&
        indexOf(toLowerCase(fileName), "20x") != -1) {

        // Open stack
        open(inputDir + fileName);

        // Remove extension
```

```
        baseName = fileName;
        baseName = replace(baseName, ".tiff", "");
        baseName = replace(baseName, ".tif", "");

        // Max projection
        run("Z Project...", "start=4 stop=24 projection=[Max Intensity]");

        // Set scale
        run("Set Scale...", "distance=2304 known=748.80 unit=microns");


        // Save MIP in MIP folder
        saveAs("Tiff", mipDir + baseName + "_MIP.tif");

        // Threshold
        setAutoThreshold("Intermodes dark");
        setOption("BlackBackground", true);
        run("Convert to Mask");

        // Save mask in Binary folder
        saveAs("Tiff", binaryDir + baseName + "_MIP_thresholded.tif");

        // Close everything safely
        while (nImages > 0) {
            close();
        }
    }
}

setBatchMode(false);

print("Done.");
```

**60x Images - ImageJ Macro:**

```
// Ask for folders
inputDir = getDirectory("Input_folder");
if (inputDir == "") exit("No input folder selected.");

outputDir = getDirectory("Output_folder");
if (outputDir == "") exit("No output folder selected.");
```

```
// Create MIP output folder
mipDir = outputDir + "MIP/";
File.makeDirectory(mipDir);

// Get file list
list = getFileList(inputDir);

setBatchMode(true);

for (i = 0; i < list.length; i++) {
   fileName = list[i];

   // Only .tif/.tiff files containing "60x"
   if ((endsWith(toLowerCase(fileName), ".tif") || endsWith(toLowerCase(fileName), ".tiff"))
&&
      indexOf(toLowerCase(fileName), "60x") != -1) {

      // Open stack
      open(inputDir + fileName);
      originalTitle = getTitle();

      // Remove extension safely
      if (endsWith(toLowerCase(fileName), ".tiff"))
         baseName = substring(fileName, 0, lengthOf(fileName) - 5);
      else
         baseName = substring(fileName, 0, lengthOf(fileName) - 4);

      // Max projection
      run("Z Project...", "start=1 stop=27 projection=[Max Intensity]");

      // Close original stack
      selectImage(originalTitle);
      close();

      // Set scale on projection
      run("Set Scale...", "distance=2304 known=249.60 unit=microns");

      // Save projection
      saveAs("Tiff", mipDir + baseName + "_MIP.tif");
```

```
        // Close projection
        close();
    }
}

setBatchMode(false);

print("Done.");
```

### SI5: CellProfiler Pipeline

We used a CellProfiler pipeline to streamline neuronal cell body analysis, developed after troubleshooting the various modules CellProfiler offers. We settled on using a combination of ImageJ and CellProfiler to process images of the *C. elegans* neuron cell bodies.

First, we created an ImageJ macro and applied it to all images, ensuring a consistent approach across the dataset. The macro was designed to receive an input .tiff file containing a Z-stack with 27 slices, consistent with our imaging parameters, and to process only files containing "20x" in the filename. First, the macro does a maximum-intensity projection (MIP) on slices 4 through 24, excluding the first and last 3 slices of the stack. Exporting our images as .tiff files from the imaging software requires rescaling the photos to microns; thus, the next step in the ImageJ pipeline was to rescale the image from pixels to microns (distance = 2304 pixels, known = 748.80 microns). We saved this image to a folder labeled MIP. We then applied a threshold to the image using the Intermodes dark algorithm and saved this binary image to a folder labeled Binary. We did this so we would have a raw MIP image to view in CellProfiler, while the binary image guides the analysis. In the CellProfiler pipeline, we can modify the neuron areas determined by the binary.

After pre-processing in ImageJ, we imported both the MIP and Binary images into CellProfiler. We used the Metadata and NamesAndTypes input modules to define how images are organized in CellProfiler. Using the Metadata tab, we applied a regular expression (RegEx) to the file names to extract the strain, treatment, and image number. Although we included a RegEx to extract similar information from the folder names, we added it as a failsafe in case there was a problem with the file-naming conventions. We pre-screened the file names to correct any discrepancies. Next, the NamesAndTypes tab ensured the MIP and Binary for the same image were correctly identified. Because both images should have the same metadata, we used this to associate them. The MIP images were labeled "Original" in this tab, while the Binary images were labeled "RawData."

Next, we added the object, measurement, and export modules to the pipeline to measure neuronal cell bodies. First, we used the "IdentifyPrimaryObjects" module to identify neurons in the images, using the "RawData" image. Because this "RawData" image was derived from the previously generated binary, we quickly processed the number of objects in the image. We set the typical object diameter range to 3 to 2000 pixels. Although we included a thresholding strategy in CellProfiler for flexibility, applying it to an already thresholded image has no effect. We used this thresholding strategy in our pipeline until we realized that thresholding preprocessing in ImageJ yielded more consistent results, since ImageJ offers more thresholding algorithms than CellProfiler. We ultimately use a thresholding algorithm in ImageJ that CellProfiler does not offer. The remaining parts of this module provide a strategy for declumping objects. We used shape as the primary method to distinguish clumped objects and propagate them, since this algorithm is intended for cases where boundaries between two objects are not always clear. We allowed the pipeline to automatically calculate the smoothing filter size for declumping and specified that local minima closer than 7 pixels should be suppressed.

Next, we used the “EditObjectsManually” module to screen the neurons in the images. This module used the “Original” image as the guiding image, with the binary overlaid onto the MIP image. In this module, we removed extraneous neurons, such as those from other worms that may have been captured unintentionally in the image, as well as any debris falsely labeled as a neuron. This module also allowed resizing areas or creating additional objects; if the binary did not align with the neurons or incorrectly merged multiple objects into a single object, we could correct it manually. Objects that were retained after this edit were labeled “Neurons_Screened.”

Following this, several image-generation modules were included to support downstream analysis. The main output image was generated using a combination of the “OverlayOutlines” module, which applied outlines of the “Neurons_Screened” objects onto the original MIP image, and the “DisplayDataOnImage” module, which added numbered labels to each outlined object so that it is clear which numbered object corresponds to which data, if necessary for additional screening. This was saved using the “SaveImages” module. We added additional image-generation modules to the pipeline, though we did not use them as final outputs. The “ConvertObjectsToImage” module overlaid the “Neurons_Screened” objects directly onto the MIP image. The “MaskImage” module applied the “Neurons_Screened” objects as a mask to the original MIP image. Although these modules were not used in the final analysis, we used them previously to aid the analysis and kept them in the pipeline for flexibility, should the outline image prove unsatisfactory.

In addition to the images, we used the “MeasureObjectIntensity” and “MeasureObjectSizeShape” modules to quantify pixel intensities and image areas within the bounds of the “Neurons_Screened” objects. The “ExportToSpreadsheet” module exports these measurements to a single Excel sheet after analyzing all images and allows selection of specific measurements for export.

### SI6: Egg Retention Raw Egg Counts

Below is additional information and statistics analysis for the egg retention assay using raw egg counts.

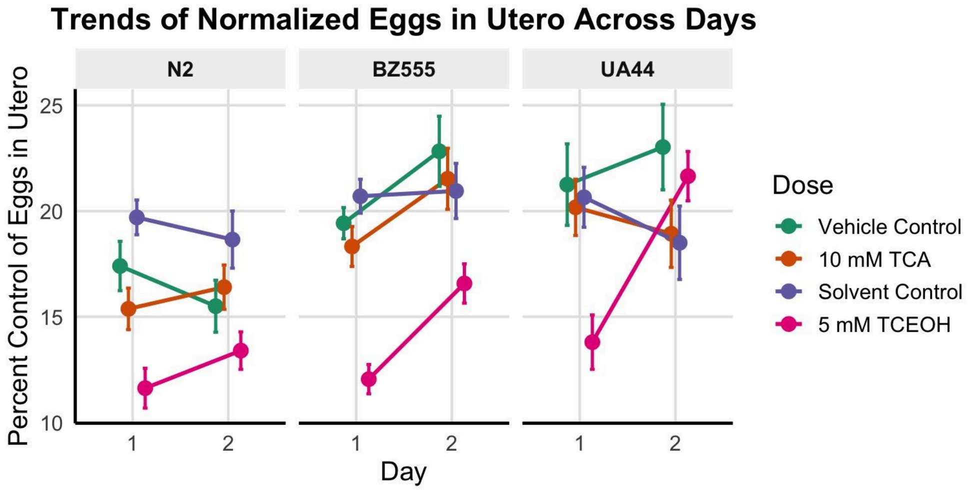


**Figure SI6.1.** Eggs in utero for N2, BZ555, and UA44 strains across Day 1 and Day 2 of adulthood. Statistics were conducted based on the control for that treatment: the vehicle control (M9) for the TCA group, and solvent control (DMSO in M9) for the TCEOH group. Three-way ANOVA for TCA- and vehicle control-treated worms indicated a significant main effect of strain ($p < 0.05$). Three-way ANOVA for TCEOH- and solvent control-treated worms indicated significant main effects of age, treatment, and strain, and significant age x treatment, and treatment x strain interactions ($p < 0.05$). The significant interactions prevented straightforward pairwise comparisons.. n = 35 - 40 worms per treatment, day, and strain across 4 experimental replicates.

| Factor | Degrees of Freedom | Sum of Squares | Mean of Squares | F-value | *p*-value |
|---|---|---|---|---|---|
| Age | 1 | 130 | 130 | 1.678 | 0.196 |
| Treatment | 1 | 252 | 252 | 3.251 | 0.072 |
| Strain | 2 | 2,183 | 1,092 | 14.067 | **< 0.001** |
| Age × Treatment | 1 | 0 | 0 | 0.004 | 0.950 |
| Age × Strain | 2 | 316 | 157.9 | 2.035 | 0.132 |
| Treatment × Strain | 2 | 86 | 43 | 0.553 | 0.576 |
| Age × Treatment × Strain | 2 | 177 | 89 | 1.142 | 0.320 |
| Residuals | 468 | 36,316 | 78 | | |

**Table SI7.1.** Results of three-way ANOVA for fecundity in TCA and vehicle control-exposed worms using raw egg counts.

| Factor | Degrees of Freedom | Sum of Squares | Mean of Squares | F-value | *p*-value |
|---|---|---|---|---|---|
| Age | 1 | 418 | 418 | 7.904 | **0.005** |
| Treatment | 1 | 3010 | 3010 | 56.896 | **< 0.001** |
| Strain | 2 | 641 | 321 | 6.059 | **0.003** |
| Age × Treatment | 1 | 975 | 975 | 18.424 | **< 0.001** |
| Age × Strain | 2 | 140 | 70 | 1.323 | 0.267 |
| Treatment × Strain | 2 | 599 | 300 | 5.661 | **0.004** |
| Age × Treatment × Strain | 2 | 288 | 144 | 2.721 | 0.067 |
| Residuals | 468 | 24,759 | 53 | | |

**Table SI7.1.** Results of three-way ANOVA for fecundity in TCEOH and solvent control-exposed worms using raw egg counts.

### SI7: Additional Statistical Analysis

Below are additional statistical analysis results, including the cumulative link mixed models used for dendrite analysis.

| Factor | Degrees of Freedom | Sum of Squares | Mean of Squares | F-value | *p*-value |
|---|---|---|---|---|---|
| Age | 1 | 10 | 10 | 0.005 | 0.943 |
| Treatment | 1 | 732 | 732 | 0.372 | 0.542 |
| Strain | 2 | 1,372 | 686 | 0.349 | 0.706 |
| Age × Treatment | 1 | 10 | 10 | 0.005 | 0.943 |
| Age × Strain | 2 | 8,152 | 4,076 | 2.072 | 0.127 |
| Treatment × Strain | 2 | 1,372 | 686 | 0.349 | 0.706 |
| Age × Treatment × Strain | 2 | 8,152 | 4,076 | 2.072 | 0.127 |
| Residuals | 468 | 920,760 | 1,967 | | |

**Table SI7.1.** Results of three-way ANOVA for fecundity in TCA and vehicle control-exposed worms.

| Factor | Degrees of Freedom | Sum of Squares | Mean of Squares | F-value | *p*-value |
|---|---|---|---|---|---|
| Age | 1 | 39,186 | 39,186 | 22.879 | **< 0.001** |
| Treatment | 1 | 59,662 | 59,662 | 34.835 | **< 0.001** |
| Strain | 2 | 29,847 | 14,923 | 8.713 | **< 0.001** |
| Age × Treatment | 1 | 39,186 | 39,186 | 22.879 | **< 0.001** |
| Age × Strain | 2 | 21,110 | 10,555 | 6.163 | **0.002** |
| Treatment × Strain | 2 | 29,847 | 14,923 | 8.713 | **< 0.001** |
| Age × Treatment × Strain | 2 | 21,110 | 10,555 | 6.163 | **0.002** |
| Residuals | 468 | 801,554 | 1713 | | |

**Table SI7.2.** Results of three-way ANOVA for fecundity in TCEOH- and solvent control-exposed worms.

| Factor | Degrees of Freedom | MSE | F-value | η²G | *p*-value |
|---|---|---|---|---|---|
| Timing | 1, 935 | 2,731 | 6.45 | 0.007 | **0.011** |
| Treatment | 1, 935 | 2,731 | 2.10 | 0.002 | 0.148 |
| Strain | 2, 935 | 2,731 | 65.09 | 0.122 | **< 0.001** |
| Timing × Treatment | 1, 935 | 2,731 | 30.86 | 0.032 | **< 0.001** |
| Timing × Strain | 2, 935 | 2,731 | 1.60 | 0.003 | 0.202 |
| Treatment × Strain | 2, 935 | 2,731 | 0.83 | 0.002 | 0.438 |
| Timing × Treatment × Strain | 2, 935 | 2,731 | 9.15 | 0.019 | **< 0.001** |

**Table SI7.3.** Results of three-way ANOVA for peristaltic speed in TCA- and vehicle-control-exposed worms.

| Factor | Degrees of Freedom | MSE | F-value | η²G | *p*-value |
|---|---|---|---|---|---|
| Timing | 1, 793 | 2,542 | 1.03 | 0.001 | 0.310 |
| Treatment | 1, 793 | 2,542 | 0.84 | 0.001 | 0.361 |
| Strain | 2, 793 | 2,542 | 21.51 | 0.051 | **< 0.001** |
| Timing × Treatment | 1, 793 | 2,542 | 0.00 | < 0.001 | 0.969 |
| Timing × Strain | 2, 793 | 2,542 | 2.47 | 0.006 | 0.085 |
| Treatment × Strain | 2, 793 | 2,542 | 0.34 | < 0.001 | 0.715 |
| Timing × Treatment × Strain | 2, 793 | 2,542 | 4.56 | 0.011 | **0.011** |

**Table SI7.4.** Results of three-way ANOVA for peristaltic speed in TCEOH- and solvent-control-exposed worms.

| **Factor** | **Degrees of Freedom** | **MSE** | **F-value** | **$\eta^2$G** | ***p*-value** |
|---|---|---|---|---|---|
| Timing | 1, 935 | 11,454 | 0.63 | < 0.001 | 0.426 |
| Treatment | 1, 935 | 11,454 | 5.34 | 0.006 | **0.021** |
| Strain | 2, 935 | 11,454 | 60.83 | 0.115 | **< 0.001** |
| Timing × Treatment | 1, 935 | 11,454 | 0.73 | < 0.001 | 0.395 |
| Timing × Strain | 2, 935 | 11,454 | 1.84 | 0.004 | 0.159 |
| Treatment × Strain | 2, 935 | 11,454 | 3.99 | 0.008 | **0.019** |
| Timing × Treatment × Strain | 2, 935 | 11,454 | 1.61 | 0.003 | 0.200 |

**Table SI7.5.** Results of three-way ANOVA for wavelength in TCA- and vehicle-control-exposed worms.

| **Factor** | **Degrees of Freedom** | **Mean of Squares** | **F-value** | **$\eta^2$G** | ***p*-value** |
|---|---|---|---|---|---|
| Timing | 1, 793 | 14,388 | 0.27 | < 0.001 | 0.605 |
| Treatment | 1, 793 | 14,388 | 19.91 | 0.024 | **< 0.001** |
| Strain | 2, 793 | 14,388 | 34.38 | 0.080 | **< 0.001** |
| Timing × Treatment | 1, 793 | 14,388 | 6.43 | 0.008 | **0.011** |
| Timing × Strain | 2, 793 | 14,388 | 10.89 | 0.027 | **< 0.001** |
| Treatment × Strain | 2, 793 | 14,388 | 3.60 | 0.009 | **0.028** |
| Timing × Treatment × Strain | 2, 793 | 14,388 | 6.28 | 0.016 | **0.002** |

**Table SI7.6.** Results of three-way ANOVA for wavelength in TCEOH- and solvent control-exposed worms.

| **Term** | **Estimate ± SE** | **Z-value** | ***p*-value** |
|---|---|---|---|
| (Intercept) | - 2.061 ± 0.401 | -5.14 | **< 0.001** |
| Strain (BZ555) | - 0.079 ± 0.504 | -0.16 | 0.876 |
| Strain (UA44) | 0.362 ± 0.480 | 0.75 | 0.451 |
| Treatment (TCA) | 0.466 ± 0.497 | 0.94 | 0.348 |
| Timing (30 min) | 0.603 ± 0.533 | 1.13 | 0.258 |
| Strain (BZ555) × Treatment | - 0.917 ± 0.721 | -1.27 | 0.203 |
| Strain (UA44) × Treatment | 0.214 ± 0.607 | 0.35 | 0.724 |
| Strain (BZ555) × Timing | - 1.235 ± 0.802 | -1.54 | 0.123 |
| Strain (UA44) × Timing | - 0.165 ± 0.655 | -0.25 | 0.802 |
| Treatment × Timing | - 1.592 ± 0.799 | -1.99 | **0.046** |
| Strain (BZ555) × Treatment × Timing | 1.665 ± 1.235 | 1.35 | 0.178 |
| Strain (UA44) × Treatment × Timing | - 0.329 ± 1.026 | -0.32 | 0.748 |

**Table SI7.7.** Results of logistic regression for the number of worms that experience omega bending behavior in TCA- and vehicle-control-treated worms. The reference levels were vehicle control for treatment, N2 for strain, and the 5-minute time point for timing.

| **Term** | **Estimate ± SE** | ***Z*-value** | ***p*-value** |
|---|---|---|---|
| (Intercept) | -2.416 ± 0.467 | -5.18 | **< 0.001** |
| Strain (BZ555) | 1.012 ± 0.554 | 1.83 | 0.068 |
| Strain (UA44) | 1.514 ± 0.518 | 2.92 | **0.003** |
| Treatment (TCEOH) | -0.086 ± 0.659 | -0.13 | 0.897 |
| Timing (30 min) | 0.401 ± 0.638 | 0.63 | 0.529 |
| Strain (BZ555) × Treatment | -0.779 ± 0.804 | -0.97 | 0.332 |
| Strain (UA44) × Treatment | -0.045 ± 0.733 | -0.06 | 0.951 |
| Strain (BZ555) × Timing | -1.461 ± 0.876 | -1.67 | 0.095 |
| Strain (UA44) × Timing | -1.371 ± 0.775 | -1.77 | 0.077 |
| Treatment × Timing | 0.099 ± 0.922 | 0.11 | 0.915 |
| Strain (BZ555) × Treatment × Timing | 1.592 ± 1.226 | 1.30 | 0.194 |
| Strain (UA44) × Treatment × Timing | -0.262 ± 1.121 | -0.23 | 0.815 |

**Table SI7.8.** Results of logistic regression for the number of worms that experience omega bending behavior in TCEOH- and solvent-control-treated worms. The reference levels were solvent control for treatment, N2 for strain, and the 5-minute time point for timing.

| Factor | Degrees of Freedom | MSE | F-value | η²G | *p*-value |
|---|---|---|---|---|---|
| Strain | 2, 12,516 | 0.002 | 11.17 | 0.002 | **< 0.001** |
| Treatment | 1, 12,516 | 0.002 | 0.84 | < 0.001 | 0.359 |
| Timing | 1, 12,516 | 0.002 | 6.04 | < 0.001 | **0.014** |
| Strain × Treatment | 2, 12,516 | 0.002 | 0.91 | < 0.001 | 0.404 |
| Strain × Timing | 2, 12,516 | 0.002 | 0.60 | < 0.001 | 0.551 |
| Treatment × Timing | 1, 12,516 | 0.002 | 0.07 | < 0.001 | 0.791 |
| Strain × Treatment × Timing | 2, 12,516 | 0.002 | 1.20 | < 0.001 | 0.301 |

**Table SI7.9.** Results of three-way ANOVA for differences in the number of high-bending angle frames across all worms in the TCA- and vehicle-control-treated worms.

| Factor | Degrees of Freedom | MSE | F-value | η²G | *p*-value |
|---|---|---|---|---|---|
| Strain | 2, 14,676 | 0.002 | 21.66 | 0.003 | **< 0.001** |
| Treatment | 1, 14,676 | 0.002 | 0.05 | < 0.001 | 0.817 |
| Timing | 1, 14,676 | 0.002 | 21.26 | 0.001 | **< 0.001** |
| Strain × Treatment | 2, 14,676 | 0.002 | 0.10 | < 0.001 | 0.904 |
| Strain × Timing | 2, 14,676 | 0.002 | 8.82 | 0.001 | **< 0.001** |
| Treatment × Timing | 1, 14,676 | 0.002 | 1.83 | < 0.001 | 0.176 |
| Strain × Treatment × Timing | 2, 14,676 | 0.002 | 1.68 | < 0.001 | 0.187 |

**Table SI7.10.** Results of three-way ANOVA for differences in the number of high-bending angle frames across all worms in the TCEOH- and solvent-control-treated worms.

| Factor | Degrees of Freedom | Sum of Squares | Mean of Squares | F-value | *p*-value |
|---|---|---|---|---|---|
| Age | 1 | 2,118 | 2,118 | 0.489 | 0.484 |
| Treatment | 1 | 9,156 | 9,156 | 2.115 | 0.146 |
| Strain | 1 | 3,367 | 3,367 | 0.778 | 0.378 |
| Age × Treatment | 1 | 2,306 | 2,306 | 0.533 | 0.466 |
| Age × Strain | 1 | 282 | 282 | 0.065 | 0.798 |
| Treatment × Strain | 1 | 3,430 | 3,430 | 0.792 | 0.374 |
| Age × Treatment × Strain | 1 | 295 | 295 | 0.068 | 0.794 |
| Residuals | 2,576 | 11,152,741 | 4,329 | | |

**Table SI7.11.** Results of three-way ANOVA for neuron cell body areas in TCA- and vehicle-control-exposed worms.

| Factor | Degrees of Freedom | Sum of Squares | Mean of Squares | F-value | *p*-value |
|---|---|---|---|---|---|
| Age | 1 | 36,568 | 36,568 | 8.919 | **0.003** |
| Treatment | 1 | 14,182 | 14,182 | 3.459 | 0.063 |
| Strain | 1 | 2,776 | 2,776 | 0.677 | 0.411 |
| Age × Treatment | 1 | 33,815 | 33,815 | 8.247 | **0.004** |
| Age × Strain | 1 | 33,058 | 33,058 | 8.063 | **0.005** |
| Treatment × Strain | 1 | 2,630 | 2,630 | 0.642 | 0.423 |
| Age × Treatment × Strain | 1 | 30,740 | 30,740 | 7.497 | **0.006** |
| Residuals | 2,469 | 10,123,205 | 4,100 | | |

**Table SI7.12.** Results of three-way ANOVA for neuron cell body areas in TCEOH- and solvent-control-exposed worms.

| Comparison | Factor | Degrees of Freedom | MSE | F-value | $\eta^2G$ | *p*-value |
|---|---|---|---|---|---|---|
| BZ555 (TCA vs. Vehicle Control) | Age | 1, 358 | 1,798 | 0.06 | < 0.001 | 0.803 |
| | Treatment | 1, 358 | 1,798 | 0.10 | < 0.001 | 0.754 |
| | Age × Treatment | 1, 358 | 1,798 | 0.01 | < 0.001 | 0.909 |
| UA44 (TCA vs. Vehicle Control) | Age | 1, 354 | 1,839 | 0.01 | < 0.001 | 0.915 |
| | Treatment | 1, 354 | 1,839 | 0.62 | 0.002 | 0.432 |
| | Age × Treatment | 1, 354 | 1,839 | 0.01 | < 0.001 | 0.907 |
| BZ555 (TCEOH vs. Solvent Control) | Age | 1, 356 | 1,956 | 3.04 | 0.008 | 0.082 |
| | Treatment | 1, 356 | 1,956 | 2.63 | 0.007 | 0.106 |
| | Age × Treatment | 1, 356 | 1,956 | 4.78 | 0.013 | **0.029** |
| UA44 (TCEOH vs. Solvent Control) | Age | 1, 354 | 1,966 | 0.04 | < 0.001 | 0.845 |
| | Treatment | 1, 354 | 1,966 | 0.22 | < 0.001 | 0.639 |
| | Age × Treatment | 1, 354 | 1,966 | 0.25 | < 0.001 | 0.617 |

**Table SI7.13.** Results of two-way ANOVA for neuron cell body areas.

| Conditions | Comparison | Degrees of Freedom | Chi-square | Raw *p*-value | Adjusted *p*-value |
|---|---|---|---|---|---|
| BZ555 × Vehicle Control | Day 1 vs. Day 3 | 2 | 4.130 | 0.127 | 1.000 |
| | Day 1 vs. Day 5 | 2 | 69.397 | **< 0.001** | **< 0.001** |
| | Day 3 vs. Day 5 | 2 | 55.590 | **< 0.001** | **< 0.001** |
| BZ555 × 10 mM TCA | Day 1 vs. Day 3 | 2 | 24.341 | **< 0.001** | **0.001** |
| | Day 1 vs. Day 5 | 2 | 59.425 | **< 0.001** | **< 0.001** |
| | Day 3 vs. Day 5 | 2 | 8.964 | **0.011** | 0.272 |
| UA44 × Vehicle Control | Day 1 vs. Day 3 | 2 | 6.476 | **0.039** | 0.942 |
| | Day 1 vs. Day 5 | 2 | 9.237 | **0.010** | 0.237 |
| | Day 3 vs. Day 5 | 2 | 0.842 | 0.657 | 1.000 |
| UA44 × 10 mM TCA | Day 1 vs. Day 3 | 2 | 3.895 | 0.143 | 1.000 |
| | Day 1 vs. Day 5 | 3 | 3.408 | 0.333 | 1.000 |
| | Day 3 vs. Day 5 | 3 | 3.013 | 0.390 | 1.000 |
| BZ555 × Day 1 | Vehicle Control vs. 10 mM TCA | 2 | 2.225 | 0.329 | 1.000 |
| BZ555 × Day 3 | | 2 | 9.001 | **0.011** | 0.267 |
| BZ555 × Day 5 | | 2 | 5.357 | 0.069 | 1.000 |
| UA44 × Day 1 | Vehicle Control vs. 10 mM TCA | 2 | 1.885 | 0.390 | 1.000 |
| UA44 × Day 3 | | 2 | 0.334 | 0.846 | 1.000 |
| UA44 × Day 5 | | 3 | 2.350 | 0.503 | 1.000 |
| Vehicle Control × Day 1 | BZ555 vs. UA44 | 2 | 58.472 | **< 0.001** | **<0.001** |
| 10 mM TCA × Day 1 | | 2 | 38.840 | **< 0.001** | **<0.001** |
| Vehicle Control × Day 3 | | 2 | 54.871 | **< 0.001** | **<0.001** |
| 10 mM TCA × Day 3 | | 2 | 85.518 | **< 0.001** | **<0.001** |
| Vehicle Control × Day 5 | | 2 | 149.429 | **< 0.001** | **<0.001** |
| 10 mM TCA × Day 5 | | 3 | 118.397 | **< 0.001** | **<0.001** |

**Table SI7.14.** Results of pairwise chi-square analysis for dendrite degeneration scores in TCA- and vehicle-control-exposed worms.

| Conditions | Comparison | Degrees of Freedom | Chi-square | Raw *p*-value | Adjusted *p*-value |
|---|---|---|---|---|---|
| BZ555 × Solvent Control | Day 1 vs. Day 3 | 4 | 18.296 | **0.001** | **0.026** |
| | Day 1 vs. Day 5 | 3 | 111.608 | **< 0.001** | **< 0.001** |
| | Day 3 vs. Day 5 | 3 | 53.761 | **< 0.001** | **< 0.001** |
| BZ555 × 5 mM TCEOH | Day 1 vs. Day 3 | 2 | 15.223 | **0.001** | **0.012** |
| | Day 1 vs. Day 5 | 2 | 80.738 | **< 0.001** | **< 0.001** |
| | Day 3 vs. Day 5 | 2 | 46.157 | **< 0.001** | **< 0.001** |
| UA44 × Solvent Control | Day 1 vs. Day 3 | 2 | 3.492 | 0.175 | 1.000 |
| | Day 1 vs. Day 5 | 2 | 14.435 | **0.001** | **0.018** |
| | Day 3 vs. Day 5 | 2 | 25.818 | **< 0.001** | **< 0.001** |
| UA44 × 5 mM TCEOH | Day 1 vs. Day 3 | 2 | 4.036 | 0.133 | 1.000 |
| | Day 1 vs. Day 5 | 2 | 10.168 | **0.006** | 0.149 |
| | Day 3 vs. Day 5 | 2 | 19.671 | **< 0.001** | **0.001** |
| BZ555 × Day 1 | Solvent Control vs. 5 mM TCEOH | 3 | 12.018 | **0.007** | 0.176 |
| BZ555 × Day 3 | | 3 | 12.068 | **0.007** | 0.172 |
| BZ555 × Day 5 | | 2 | 15.965 | **< 0.001** | **0.008** |
| UA44 × Day 1 | Solvent Control vs. 5 mM TCEOH | 2 | 1.459 | 0.482 | 1.000 |
| UA44 × Day 3 | | 2 | 0.979 | 0.613 | 1.000 |
| UA44 × Day 5 | | 2 | 5.151 | 0.076 | 1.000 |
| Solvent Control × Day 1 | BZ555 vs. UA44 | 3 | 9.923 | **0.019** | 0.462 |
| 5 mM TCEOH × Day 1 | | 2 | 0.189 | 0.910 | 1.000 |
| Solvent Control × Day 3 | | 3 | 47.121 | **< 0.001** | **< 0.001** |
| 5 mM TCEOH × Day 3 | | 2 | 30.820 | **< 0.001** | **< 0.001** |
| Solvent Control × Day 5 | | 2 | 78.833 | **< 0.001** | **< 0.001** |
| 5 mM TCEOH × Day 5 | | 2 | 53.662 | **< 0.001** | **< 0.001** |

**Table SI7.15.** Results of pairwise chi-square analysis for dendrite degeneration scores in TCEOH- and solvent-control-exposed worms.

| Term | Estimate ± SE | Z-value | *p*-value |
|---|---|---|---|
| Strain (UA44) | -3.202 ± 0.496 | -6.454 | **< 0.001** |
| Age (Day 3) | 0.750 ± 0.477 | 1.574 | 0.116 |
| Age (Day 5) | 4.661 ± 0.640 | 7.283 | **< 0.001** |
| Treatment (TCA) | -0.214 ± 0.473 | -0.453 | 0.651 |
| Strain (UA44) × Age (Day 3) | 0.196 ± 0.664 | 0.296 | 0.768 |
| Strain (UA44) × Age (Day 5) | -3.498 ± 0.778 | -4.495 | **< 0.001** |
| Strain (UA44) × Treatment | 0.609 ± 0.663 | 0.919 | 0.358 |
| Age (Day 3) × Treatment | 1.406 ± 0.693 | 2.030 | **0.042** |
| Age (Day 5) × Treatment | -0.998 ± 0.791 | -1.263 | 0.207 |
| Strain (UA44) × Age (Day 3) × Treatment | -1.751 ± 0.956 | -1.832 | 0.067 |
| Strain (UA44) × Age (Day 5) × Treatment | 0.412 ± 1.031 | 0.400 | 0.689 |

**Table SI7.16.** Results of cumulative link mixed-effects modeling for dendrite degeneration scores in TCA- and vehicle-control-exposed worms. The reference levels were vehicle control for treatment, BZ555 for strain, and day 1 of adulthood for age.

| Term | Estimate ± SE | Z-value | *p*-value |
|---|---|---|---|
| Strain (UA44) | -1.398 ± 0.511 | -2.736 | **0.006** |
| Age (Day 3) | 1.187 ± 0.510 | 2.327 | **0.020** |
| Age (Day 5) | 4.865 ± 0.610 | 7.970 | **< 0.001** |
| Treatment (TCEOH) | -1.520 ± 0.526 | -2.889 | **0.004** |
| Strain (UA44) × Age (Day 3) | -1.759 ± 0.732 | -2.404 | **0.016** |
| Strain (UA44) × Age (Day 5) | -3.419 ± 0.775 | -4.411 | **< 0.001** |
| Strain (UA44) × Treatment | 1.214 ± 0.734 | 1.653 | 0.098 |
| Age (Day 3) × Treatment | 0.429 ± 0.727 | 0.590 | 0.555 |
| Age (Day 5) × Treatment | -0.220 ± 0.759 | -0.290 | 0.772 |
| Strain (UA44) × Age (Day 3) × Treatment | -0.445 ± 1.043 | -0.427 | 0.670 |
| Strain (UA44) × Age (Day 5) × Treatment | 0.138 ± 1.053 | 0.131 | 0.896 |

**Table SI7.17.** Results of cumulative link mixed-effects modeling for dendrite degeneration scores in TCEOH- and solvent-control-exposed worms.

| **Conditions** | **Comparison** | **Day 1 EMM ± SE** | **Day 3 EMM ± SE** | **Day 5 EMM ± SE** | **Z-ratio** | **Adjusted *p*-value** |
|---|---|---|---|---|---|---|
| BZ555 × Vehicle Control | Day 1 - Day 3 | -2.665 ± 0.493 | -1.915 ± 0.486 | 1.997 ± 0.544 | -1.574 | 0.257 |
| | Day 1 - Day 5 | | | | -7.283 | **< 0.001** |
| | Day 3 - Day 5 | | | | -6.279 | **< 0.001** |
| BZ555 × 10 mM TCA | Day 1 - Day 3 | -2.878 ± 0.490 | -0.722 ± 0.514 | 0.785 ± 0.536 | -4.189 | **< 0.001** |
| | Day 1 - Day 5 | | | | -6.502 | **< 0.001** |
| | Day 3 - Day 5 | | | | -2.706 | **0.019** |
| BZ555 × Solvent Control | Day 1 - Day 3 | -5.283 ± 0.540 | -4.097 ± 0.525 | -0.419 ± 0.457 | -2.327 | 0.052 |
| | Day 1 - Day 5 | | | | -7.970 | **< 0.001** |
| | Day 3 - Day 5 | | | | -6.237 | **< 0.001** |
| BZ555 × 5 mM TCEOH | Day 1 - Day 3 | -6.803 ± 0.601 | -5.187 ± 0.549 | -2.158 ± 0.505 | -3.059 | **0.006** |
| | Day 1 - Day 5 | | | | -7.751 | **< 0.001** |
| | Day 3 - Day 5 | | | | -5.471 | **< 0.001** |
| UA44 × Vehicle Control | Day 1 - Day 3 | -5.866 ± 0.529 | -4.920 ± 0.516 | -4.702 ± 0.517 | -2.024 | 0.106 |
| | Day 1 - Day 5 | | | | -2.461 | **0.037** |
| | Day 3 - Day 5 | | | | -0.456 | 0.892 |
| UA44 × 10 mM TCA | Day 1 - Day 3 | -5.471 ± 0.524 | -4.869 ± 0.507 | -4.893 ± 0.515 | -1.303 | 0.394 |
| | Day 1 - Day 5 | | | | -1.232 | 0.434 |
| | Day 3 - Day 5 | | | | 0.052 | 0.999 |
| UA44 × Solvent Control | Day 1 - Day 3 | -6.682 ± 0.586 | -7.254 ± 0.615 | -5.236 ± 0.557 | 1.099 | 0.515 |
| | Day 1 - Day 5 | | | | -2.797 | **0.014** |
| | Day 3 - Day 5 | | | | -3.787 | **< 0.001** |
| UA44 × 5 mM TCEOH | Day 1 - Day 3 | -6.988 ± 0.599 | -7.576 ± 0.635 | -5.625 ± 0.563 | 1.091 | 0.520 |
| | Day 1 - Day 5 | | | | -2.593 | **0.026** |
| | Day 3 - Day 5 | | | | -3.549 | **0.001** |

**Table SI7.18.** Effect of age within strain and dose for the dendrite degeneration assay using latent estimated marginal means from the cumulative link mixed-effects modeling. Given that for each strain and dose there were 3 comparisons (day 1 vs. day 3, day 1 vs. day 5, and day 3 vs. day 5), *p*-values were adjusted for multiple comparisons using Tukey's method. Latent EMMs are reported in this table.

| Conditions | Comparison | Estimate (Difference) ± SE | Z-ratio | *p*-value |
|---|---|---|---|---|
| Vehicle Control × Day 1 | BZ555 - UA44 | 3.202 ± 0.496 | 6.454 | **< 0.001** |
| Vehicle Control × Day 3 | | 3.005 ± 0.498 | 6.033 | **< 0.001** |
| Vehicle Control × Day 5 | | 6.699 ± 0.684 | 9.792 | **< 0.001** |
| 10 mM TCA × Day 1 | | 2.592 ± 0.486 | 5.333 | **< 0.001** |
| 10 mM TCA × Day 3 | | 4.147 ± 0.542 | 7.651 | **< 0.001** |
| 10 mM TCA × Day 5 | | 5.678 ± 0.608 | 9.332 | **< 0.001** |
| Solvent Control × Day 1 | | 1.398 ± 0.511 | 2.736 | **0.006** |
| Solvent Control × Day 3 | | 3.157 ± 0.550 | 5.746 | **< 0.001** |
| Solvent Control × Day 5 | | 4.818 ± 0.623 | 7.735 | **< 0.001** |
| 5 mM TCEOH × Day 1 | | 0.185 ± 0.529 | 0.349 | 0.727 |
| 5 mM TCEOH × Day 3 | | 2.388 ± 0.547 | 4.370 | **< 0.001** |
| 5 mM TCEOH × Day 5 | | 3.466 ± 0.566 | 6.122 | **< 0.001** |

**Table SI7.19.** Effect of strain within day and dose for the dendrite degeneration assay using latent estimated marginal means from the cumulative link mixed-effects modeling. Because each strain and dose had only one comparison (BZ555 vs. UA44), *p*-values were not adjusted for multiple comparisons. Latent EMMs are reported in this table.

| **Strain** | **Comparison** | **Control EMM ± SE** | **Treatment EMM ± SE** | **Z-value** | ***p*-value** |
|---|---|---|---|---|---|
| BZ555 × Day 1 | Vehicle Control - TCA | - 2.665 ± 0.493 | - 2.878 ± 0.490 | 0.453 | 0.651 |
| BZ555 × Day 3 | | - 1.915 ± 0.486 | - 0.722 ± 0.514 | - 2.361 | **0.018** |
| BZ555 × Day 5 | | 1.997 ± 0.544 | 0.785 ± 0.536 | 1.910 | 0.056 |
| BZ555 × Day 1 | Solvent Control - TCEOH | - 5.283 ± 0.540 | - 6.803 ± 0.601 | 2.889 | **0.004** |
| BZ555 × Day 3 | | - 4.097 ± 0.525 | - 5.187 ± 0.549 | 2.129 | **0.033** |
| BZ555 × Day 5 | | - 0.419 ± 0.457 | - 2.158 ± 0.505 | 3.084 | **0.002** |
| UA44 × Day 1 | Vehicle Control - TCA | - 5.866 ± 0.529 | - 5.471 ± 0.524 | - 0.852 | 0.394 |
| UA44 × Day 3 | | - 4.920 ± 0.516 | - 4.869 ± 0.507 | - 0.110 | 0.913 |
| UA44 × Day 5 | | - 4.702 ± 0.517 | - 4.893 ± 0.515 | 0.402 | 0.688 |
| UA44 × Day 1 | Solvent Control - TCEOH | - 6.682 ± 0.586 | - 6.988 ± 0.599 | 0.595 | 0.552 |
| UA44 × Day 3 | | - 7.254 ± 0.615 | - 7.576 ± 0.635 | 0.594 | 0.553 |
| UA44 × Day 5 | | - 5.236 ± 0.557 | - 5.625 ± 0.563 | 0.750 | 0.453 |

**Table SI7.20.** Effect of dose within age and strain for the dendrite degeneration assay using latent estimated marginal means from the cumulative link mixed-effects modeling. Because each strain and dose had only one comparison (control vs. treatment), *p*-values were not adjusted for multiple comparisons. Latent EMMs are reported in this table.

### SI8: Additional Information for Assessing Exposure

MZmine (Version 4.8.0)was used to quickly assess detection of TCA in the worms and to find the presence of chlorinated species in the worms treated with TCEOH. As the instrumental method we used is only able to detect trichloroethanol glucuronide (TCEOH-gluc) rather than trichloroethanol, and the worms appear to be unable to metabolize TCEOH to TCEOH-gluc, we looked for the presence of chlorinated species to justify that worms had an internal dose of TCEOH. The worms for this analysis were dosed in a similar fashion to the worms in the main text, but they were solely collected for metabolomics. Note that quantitation was not possible for these compounds.

Below are the inputs that were used for MZmine analysis using the MZwizard tool:

Setup:

UHPLC - Orbitrap - DDA

UHPLC Module:

Check "Smoothing"
Check "Stable ionization across samples"
Crop retention time: 0.30 - 15 min
Max peaks in chromatogram: 15
Minimum consecutive scans: 4
Approximate feature FWHM: 0.05 minutes
RT tolerance (intra-sample): 0.04 minutes
RT tolerance (sample-to-sample): 0.10 minutes

Orbitrap Module:

Select appropriate ion mode (negative or positive); Factor of lowest signal
Noise threshold MS1: 5.00
Noise threshold MS2…MSn: 2.50
Minimum feature height: 1.0E4
m/z tolerance (scan-to-scan): 0.0020 m/z or 10.0 ppm
m/z tolerance (intra-sample): 0.0015 m/z or 3.0 ppm
m/z tolerance (sample-to-sample): 0.0015 m/z or 5.0 ppm

Filters Module:

Original feature list: REMOVE
Minimum aligned samples: At least 5 samples AND 0.0%
Minimum aligned samples (in any group): At least 5 samples AND 0.0% in ANY group in condition (strain)
Uncheck "Pooled QC area RSD ≤ 20%
Check "Only keep features with 13C"
Uncheck "Apply strict shape filtering"

Annotation Module:

Check “Annotate lipids” and import desired annotation libraries

DDA Module:

Check “Apply spectral networking (FBMN/IIMN)”; other functions in this module can be done after initial analysis to improve analysis runtime.

After preparing the MZwizard, navigate to “Create batch.” Add the following step to the batch:

Feature list blank subtraction:

Aligned feature list: Those created by previous batch step
Blank/Control raw data files: Specific raw data files (select your blanks)
Minimum # of detection in blanks: 7
Quantification: Area
Ratio type: MAXIMUM
Fold change increase: 300.0 %
Keep or remove features (of rows) below fold change: REMOVE - Only keep features above fold change
Check “Create secondary list of subtracted features
Suffix: subtracted

Below are boxplots of exposure for varying compounds, which were generated using RStudio. Analysis was split between the TCA and vehicle control-treated worms, and the TCEOH and solvent control-treated worms. The mass spectrum of TCA exhibits prominent peaks at *m/z* 160.8970, 162.8940, 164.8910, and 166.8881 in an isotopic pattern of 27:27:9:1 (M, M + 2, M + 4, and M + 6) that is characteristic of a trichlorinated species. The mass spectrum of the likely chlorinated species that seems to be more present in the TCEOH treated worms exhibits prominent peaks at *m/z* 388.9370, 390.9341, 392.9309, and 394.9282 in an isotopic pattern of 27:27:9:1 (M, M + 2, M + 4, and M + 6) that is characteristic of a trichlorinated species.

We see that for TCA, the compound is largely detected among the TCA-treated worms and largely not detected among the vehicle control-treated worms (Figure S18.1). In comparison, TCA is generally not detected in the TCEOH- and solvent control-treated worms or it is detected at low levels (Figure S18.2). Overall, these trends followed what was expected based on the exposures of the worms, in that we expected the TCA-treated worms to have the greatest detection of TCA. Instrument carryover may be responsible for the detection of TCA among the other treatments. It is also possible that a larger sample size may improve the detection and stabilize the estimates we provide here today.

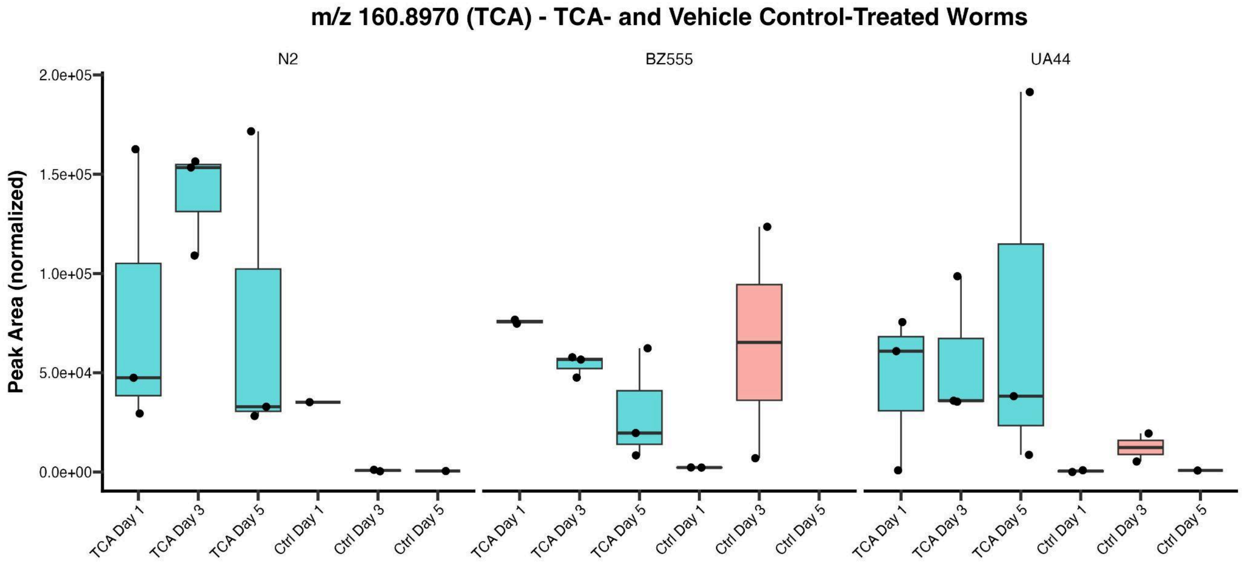


**Figure SI8.1.** Boxplot of TCA peak areas across treatments for TCA and vehicle-control-treated worms. Treatments with empty boxplots or lines are indicative of there being no detection of TCA or some samples with no detection. N = 3 replicates per treatment.

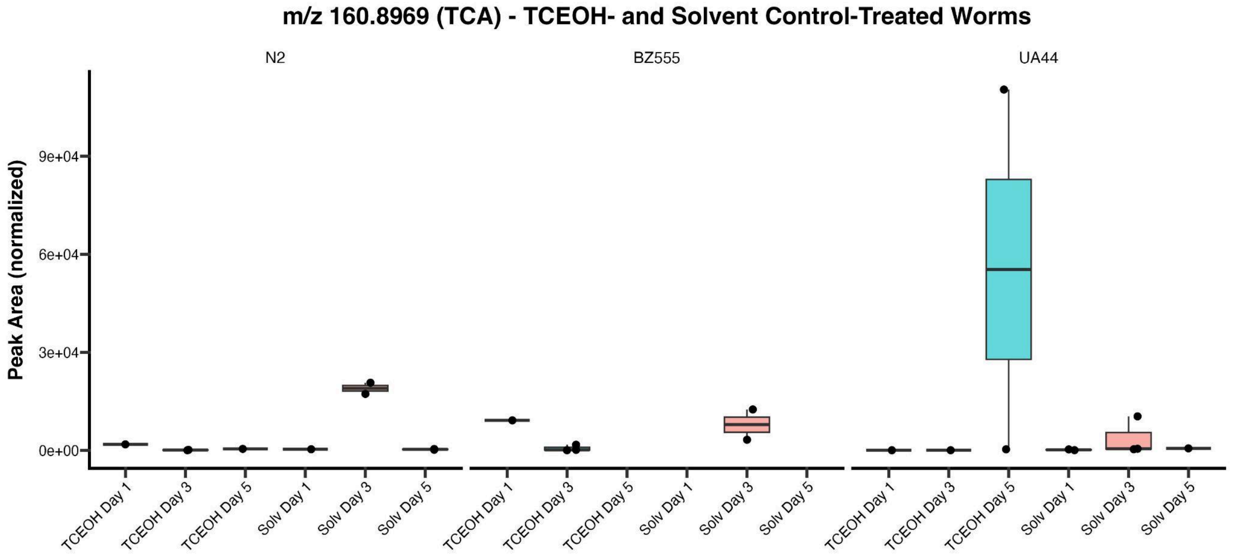


**Figure SI8.2.** Boxplot of TCA peak areas across treatments for TCEOH and solvent-control-treated worms. Treatments with empty boxplots or lines are indicative of there being no detection of TCA or some samples with no detection. N = 3 replicates per treatment.

On average, the unknown chlorinated compound is largely not detected among the TCA and vehicle control-treated worms (Figure S18.3). Comparatively, the compound is detected regularly in the TCEOH-treated worms, and generally not detected in the solvent control-treated worms or detected at low levels (Figure S18.4). Overall, these trends followed what was expected based on the exposures of the worms, though it is important to note that it would have been ideal to be able to detect TCEOH directly. Without knowing the identity of the compound, it is hard to assess what this compound is in relation to TCEOH. Using SIRIUS, which can be used for elucidating chemical structures from mass spectrum, we found that the most likely molecular formula for this unknown compound is $C_8H_{14}Cl_3O_9P$ (Böcker et al. 2009). Given its minimal presence in the TCA-treated worms, this compound is likely a TCEOH metabolite, though its detection is not as stable as that of TCA even after considering the small sample size. Again, it is possible that its detection in other treatments could be the result of instrument carryover.

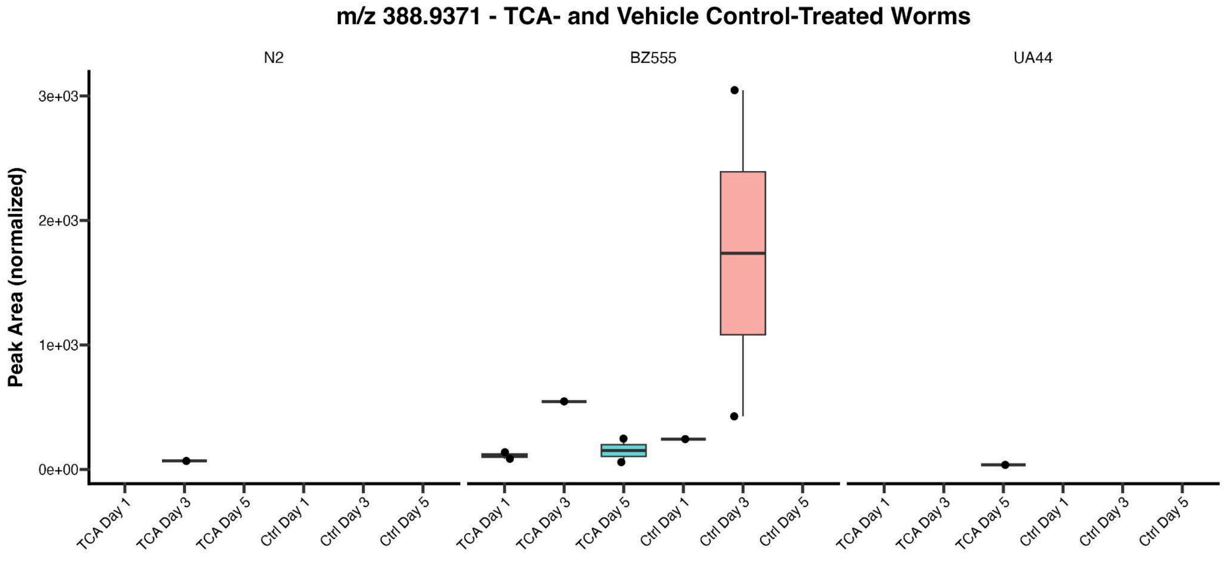


**Figure SI8.3.** Boxplot of unknown chlorinated species peak areas across treatments for TCA and vehicle-control-treated worms. Treatments with empty boxplots or lines are indicative of there being no detection of the compound or some samples with no detection. N = 3 replicates per treatment.

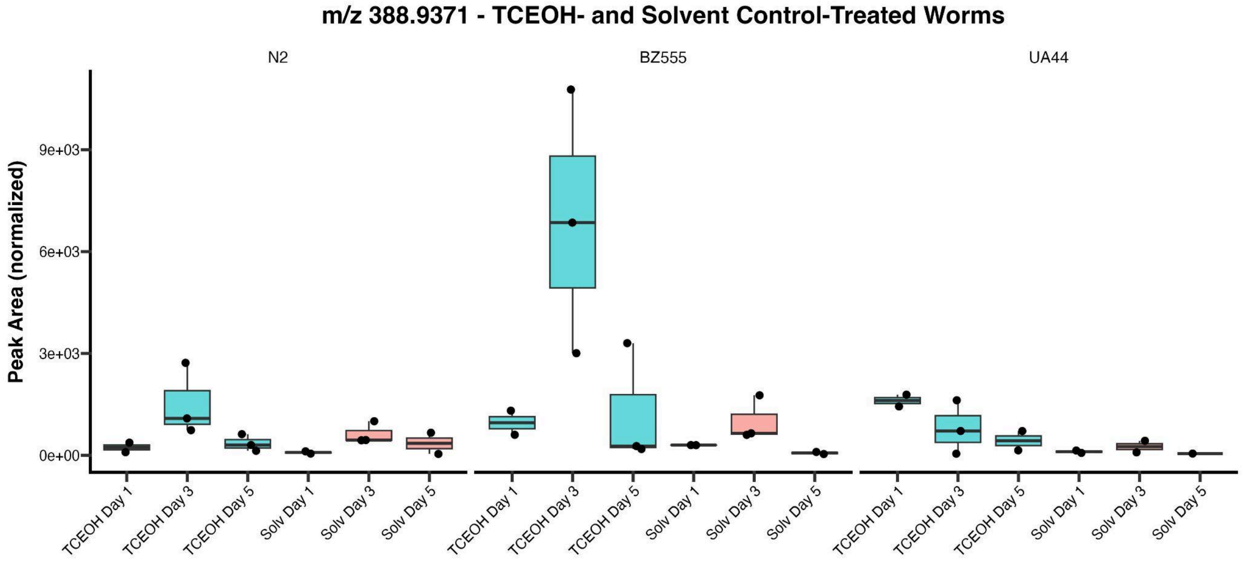


**Figure SI8.4.** Boxplot of unknown chlorinated species peak areas across treatments for TCEOH and solvent-control-treated worms. Treatments with empty boxplots or lines are indicative of there being no detection of the compound or some samples with no detection. N = 3 replicates per treatment.
